# Genome - nuclear lamina interactions are multivalent and cooperative

**DOI:** 10.1101/2024.01.10.574825

**Authors:** Lise Dauban, Mathias Eder, Marcel de Haas, Vinícius H. Franceschini-Santos, J. Omar Yañez-Cuna, Tom van Schaik, Christ Leemans, Koen Rademaker, Miguel Martinez Ara, Moreno Martinovic, Elzo de Wit, Bas van Steensel

## Abstract

Lamina-associated domains (LADs) are megabase-sized genomic regions that interact with the nuclear lamina (NL). It is not yet understood how their interactions with the NL are encoded in their DNA. Here, we designed an efficient LAD “scrambling” approach, based on transposon-mediated local hopping of loxP recombination sites, to generate series of large deletions and inversions that span LADs and flanking sequences. Mapping of NL interactions in these rearrangements revealed that a single LAD contacts the NL through multiple regions that act cooperatively or redundantly; some have more affinity for the NL than others and can pull neighbouring sequences to the NL. Genes drawn towards the NL showed often, but not always, reduced expression and increased H3K9me3 levels. Furthermore, neighbouring LADs can cooperatively interact with the NL when placed close enough to each other. These results elucidate principles that govern the positioning of megabase-sized genomic regions inside the cell nucleus.

**Highlights:**

- Efficient generation of LAD rearrangements by transposon hopping of *loxP* elements.
- NL interactions are multivalent, with some subregions being more potent tethers than others.
- LADs can cooperate to promote their association to the NL.
- Changes in NL association are often accompanied by changes in gene activity and H3K9me3.

## INTRODUCTION

*LADs are major architectural regions of the genome.* The intricate folding of the genome inside a cell nucleus is controlled at multiple levels ^1,2^. In metazoan nuclei, one of the key mechanisms is thought to involve contacts of the genome with the nuclear lamina (NL), a proteinaceous meshwork that coats the inner nuclear membrane. These contacts are mediated by large genomic regions named lamina-associated domains (LADs) ^2-5^. In most human and mouse cell types there are about 1,000 of these LADs, each spanning 0.1 - 10 Mb and collectively covering about 40% of the genome ^6-9^. Polymer simulations, in some cases supported by experimental evidence, indicate that LAD-NL interactions play a key role in the spatial organisation of chromosomes ^10-14^.

*Role of NL interactions in gene repression.* In addition to a probable role in chromosome folding, NL interactions exhibit an intriguing inverse relationship with local gene activity. Genes within LADs are mostly transcriptionally inactive ^6,7,15-17^. Possibly, NL contacts play a causal role in gene silencing. This notion is supported by studies in which forced NL anchoring of a locus (by means of a LacO array that was bound by a NL-tethered LacI protein) caused reduced transcription of some, but not all, genes around the anchoring site ^18,19^. However, another study using a highly similar approach found no evidence for gene repression ^20^. Because of the artificial nature of the tethering mechanism that was used in each of these studies, it is still unknown how natural NL-tethering mechanisms may contribute to gene repression.

*How are LAD-NL interactions instructed by LAD DNA*? Although several chromatin and NL proteins have been identified that may mediate NL-LAD interactions ^21,22^, a major unresolved question is how these interactions are encoded within LAD DNA. This is particularly intriguing considering the enormous size of LADs. For example, a LAD of 3 Mb consists of ∼1 mm of linear DNA, or roughly 150 µm of nucleosomal 10 nm fibre. How is the anchoring of such a giant molecule to the NL instructed by its own DNA sequence?

*Possible principles of LAD anchoring*. Single-cell DamID maps indicate that LAD DNA contact the NL in continuous stretches ^23^. Several models may explain these broad interactions. First, LADs may have affinity for the NL along their entire length. This could be due to a dense peppering of LAD DNA with sequence elements that mediate NL interactions; or to the near-uniform coating of LADs by a histone modification or NL-binding chromatin protein. Alternatively, each LAD may contain only one or a few tethering elements. The remainder of the LAD may then be secured to the NL by a zipper-like mechanism that spreads from the initial tethering site, or passively contact the NL by Brownian motion ^24,25^. Hybrids between these two main classes of models are also possible: LADs could contain one or more tethering elements that drive NL contacts with relatively high affinity, complemented by a multitude of elements that interact only weakly with the NL.

*Candidate NL anchoring elements.* Three previous reports have described small-scale screens to identify NL-tethering elements, in which the localisation of randomly integrated LacO repeats fused to LAD-derived DNA sequences was assessed by fluorescence microscopy ^26-28^. Multiple LAD fragments, some only a few kb in size, were identified that appeared to have the potential to promote NL interactions. However, in most experiments these sequences were integrated randomly, in unknown genomic locations. This hampers interpretation of the results, because the ability of some fragments to target to the NL was demonstrated to be context-dependent ^27^. Moreover, homing effects ^29,30^ could have biased the integrations towards specific chromatin contexts. Critically, the role of the candidate LAD-derived elements was not further tested in their native LAD context. Another limitation of these early studies was that they relied strictly on fluorescence microscopy of tagged loci, which was unable to reveal the detailed pattern of NL contacts in the tested loci. Systematic deletions and rearrangements of LAD DNA in its natural location, together with high-resolution mapping of changes in NL contact across the entire locus, would provide valuable insights into the mechanisms that drive LAD - NL interactions.

*Possible synergy between neighbouring LADs*. Besides this, the interplay between neighbouring LADs is also poorly understood. Possibly, neighbouring LADs interact with the NL in a coordinated manner rather than independently. LADs contact each other in 3D space ^31,32^. Moreover, single-cell maps revealed coordinated NL interactions of neighbouring LADs on the same chromosome ^23^. However, beyond such correlative data, there is no perturbation-based evidence that neighbouring LADs may influence each other’s NL interaction, across the intervening inter-LAD (iLAD).

*Aim of this study and experimental approach.* Here, we report the systematic dissection of LADs in mouse cells, addressing several of the unresolved issues highlighted above, such as the principles of LAD-NL contacts, LAD-LAD cooperativity and the link between NL interaction and gene repression. To study LADs in their native context, we developed an efficient technology to create series of deletions and inversions of up to ∼2 Mb. This technology combines local “hopping” of the *Sleeping Beauty* transposon (SB) with Cre-mediated recombination between two *loxP* sites ^33-35^.

*Brief description of the technique.* SB is a transposable element that follows a cut-and- paste transposition mechanism. Upon transposition, SB is excised and generally re-integrates nearby on the same chromosome, within ∼2 Mb from its original location ^33^. Moreover, mapping genomic integrations after SB hopping from a plasmid setting reported that these integrations are not biased towards any chromatin type ^36-39^. In the technique employed here, one *loxP* site is inserted in a fixed position in the genome, while the other is being “hopped” around in the same locus by SB transposition. Next, by Cre-mediated recombination of the resulting *loxP* pairs, we generate large deletions and inversions throughout a given locus, enabling us to effectively perturb the long-range sequence arrangement of LADs and study the effects on NL interactions and gene regulation.

*Summary of results.* Using this technology, we dissected a ∼10 Mb region comprising four LADs. We generated a panel of large deletions and inversions with high efficiency and mapped the effects on NL interactions, gene expression and heterochromatin marks in detail. Our data show that LAD anchoring is multivalent, with some tethering elements being more potent than others. These elements can associate with the NL autonomously to various degrees, and can drag up to ∼200 kb of neighbouring iLAD sequences towards the NL. Genes in the affected iLAD regions are often repressed, and we observe spreading of heterochromatin. We also show that neighbouring LADs can cooperate to promote their association with the NL. Together, our results advance the understanding of the mechanisms of LAD - NL interactions and the functional consequences.

## RESULTS

### Experimental design

*Overview of the method*. To identify regions in LADs that may mediate NL interactions, we adapted a transposon-based strategy ^33-35^ to generate large deletions and inversions in LADs and the flanking iLAD regions. This technique consists of three main steps. First, a cassette consisting of two *loxP* recombination sites, one of which is embedded in a SB transposon, is integrated into a genomic locus of interest. Second, SB hopping is induced by transient expression of SB transposase: by a cut-and-paste mechanism, the transposon is excised and typically re-integrates semi-randomly within 2 Mb from its original site (**Figure 1A, left**) ^33,36,40^. Thus, one *loxP* (the anchor *loxP*) remains in the original insertion position, while the other (the hopped *loxP*) is relocated to various positions in the locus. Third, recombination between the two *loxP* sites is induced by expression of Cre recombinase (**Figure 1A, right**). The outcome of the recombination reaction depends on the orientation of the hopped *loxP* relative to the anchor *loxP*: if the *loxP* sites are in the same orientation, the sequences in between them will be excised from the genome, leading to a deletion; if the *loxP* sites are in the opposite orientation, the sequences will be flipped, leading to an inversion. This powerful approach allows the generation of clonal cell lines with a wide diversity of inversions and deletions in a region of interest.

**Figure 1:**
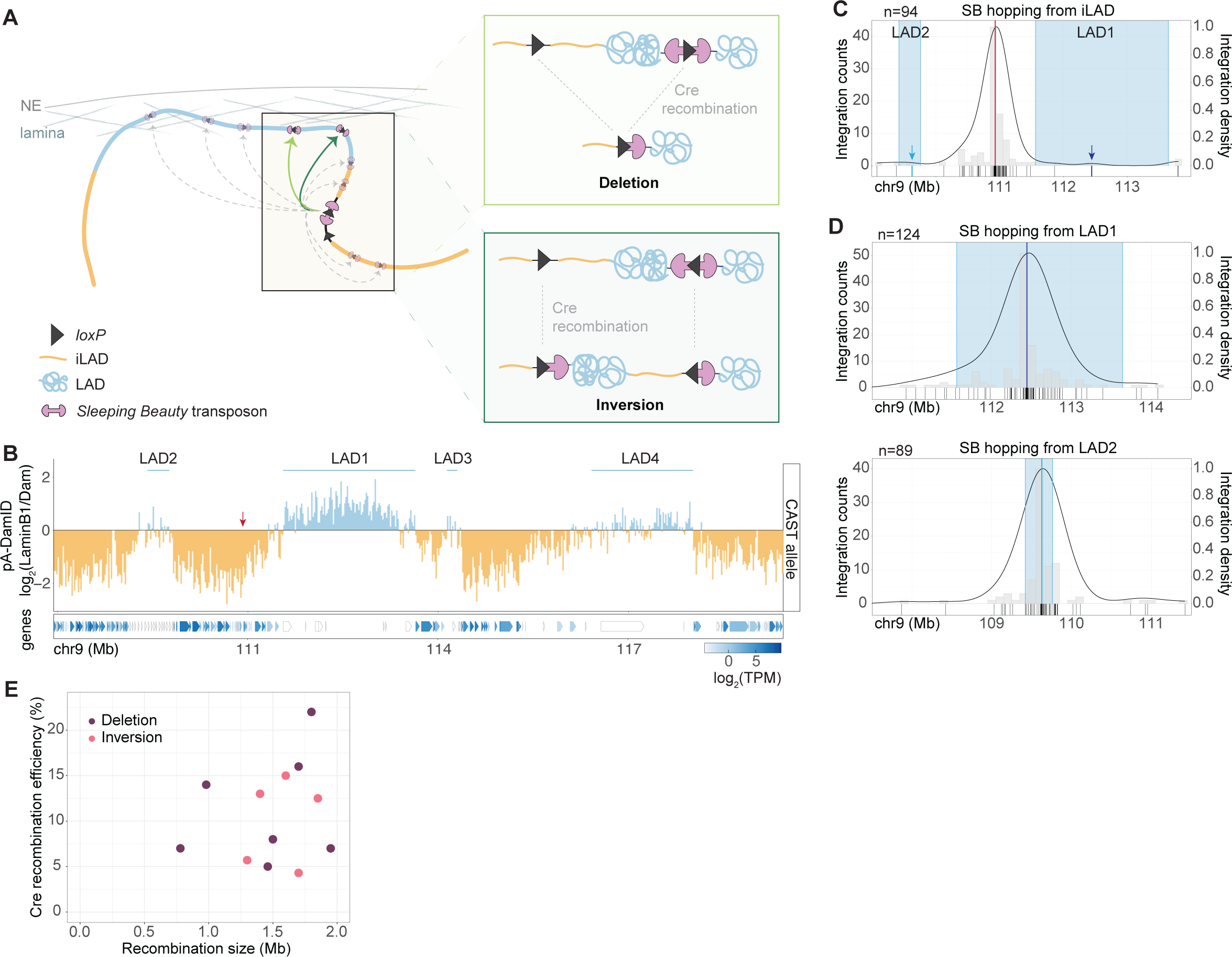
Strategy to generate large inversions and deletions that disrupt LADs. **(A)** Left panel: A cassette containing two *loxP* recombination sequences, one of which is embedded within a SB transposon, is inserted in an iLAD (orange). After SB hopping, clones with SB integrations in the nearby LAD are selected. In these clones, one *loxP* is now located inside the LAD, while the other *loxP* is still at the original iLAD position. NE: nuclear envelope. Right panel: Cre-mediated recombination generates a deletion (top) or an inversion (bottom), depending on the relative orientation of the two *loxP* sites. **(B)** Top: CAST allele-specific pA-DamID track of NL interactions, across the locus of interest on chromosome 9, in mESCs. Bin size is 20 kb. The *loxP*-SB cassette is inserted in the LAD1-flanking iLAD (arrow). iLADs are coloured in orange; LADs in blue. Blue horizontal lines mark four LADs. Genes in the bottom track are coloured according to their expression level in wild-type (WT) cells (not allele-specific). **(C)** Distribution of SB integrations after hopping from the launch site, shown as a histogram of the number SB integrations per bin of 100 kb (left y-axis) as well as a density curve (right y-axis). Each tick on the x-axis represents a unique SB integration event. SB integrations were mapped both in cell pools and clonal cell lines; n is the number of unique integrations in the plotted window. Red line: launch site, blue rectangles: LADs. Blue arrows mark SB integrations used in subsequent re-hopping experiments in panel D. **(D)** Same as in (C) for SB hopping from LAD1 (top) or LAD2 (bottom). **(E)** Cre-recombination efficiency appears largely independent of the distance between the two loxP sites, for the 12 recombinations reported in this study.

*Improvements over previous methodology*. Compared to previous work that combined SB hopping and *loxP* recombination to generate genomic rearrangements ^33,34^, we implemented several improvements. First, we did not integrate any selectable marker gene together with the *loxP*-SB cassette, because such a gene is likely to affect the LAD architecture^41^. Second, we used an enhanced version of the SB transposase (SB100x), which is 10-100 times more active than its previous version SB11 ^42^. Third, mapping of SB integrations was also improved by the use of the TagMap technique ^43^, which is substantially more scalable than other methods. Finally, we selected Cre recombinase-transfected cells, enabling a more efficient screening for recombined clones. We thus improved every step of the technique, leading to a highly efficient generation and detection of hopped and recombined clones.

*Model cell line.* As a model system, we chose mouse embryonic stem cells (mESCs), which have a well-characterized LAD architecture ^7,9,32,44^. We conducted all experiments in F1 hybrid (129/Sv:CAST/EiJ) mESCs, in which the CAST and the 129S1 alleles can be easily distinguished by sequencing-based methods due to the high density of sequence polymorphisms. The benefit of this is that we could monitor effects of inversions and deletions in one allele, while the other, unperturbed allele served as a control in the same cells. An additional advantage of this is approach is that heterozygous perturbations are less likely to compromise cellular homeostasis or differentiation state than homozygous ones.

### Efficient generation of genomic rearrangements involving LADs

*Locus of interest.* For this study we chose a ∼10 Mb region on chromosome 9, harbouring four LADs (here named LAD1 through LAD4) of varying length and interaction frequency with the NL (**Figure 1B**). The locus includes dozens of genes, enabling us to link any effects of the rearrangements on NL interactions to changes in gene expression.

*Hopping is mostly confined to spatial compartments.* We integrated the *loxP*-SB cassette in the CAST allele of mESCs, in an intergenic iLAD sequence, ∼630 kb away from LAD1 (**Figure 1B, arrow**). We then induced SB hopping by transient expression of the SB transposase, isolated clonal cell lines, screened these for SB hopping from the launch site by PCR (**Figure S1A**), and finally mapped the new integration sites by TagMap ^43^. Out of a total of 183 screened clones, this yielded 30 clones with precisely mapped hopping events. We also mapped SB integrations in pools of cells. The combined data showed that most SB integrations remained confined to a region of ∼2 Mb around the launch site (**Figure 1C**); out of 142 integrations found on chromosome 9, 88 were located less than 1 Mb away from the launch site. This is in line with earlier studies ^33,36,40^. Surprisingly, we detected only three hopping events within the nearby LADs. To test whether this apparent constraint on SB hopping was specific for the selected locus, we inserted the same *loxP*-SB cassette on other chromosomes, either closer to a LAD border (300 kb) (**Figure S1B**) or in a smaller inter-LAD surrounded by two large LADs (**Figure S1C**). Again, we found that most hopped SB integrations remained in iLADs (324/341 and 205/226), and were even strongly skewed away from the nearby LAD in one of the two locations (**Figure S1B**). These results suggested either that LAD chromatin is refractory to SB integration or that hopping from an iLAD to a LAD is less efficient than within iLADs. To test these two hypotheses, we re-induced SB hopping in two clones in which SB had integrated within LAD1 or LAD2 (**Figure 1D**). Mapping of SB integration sites in the resulting re-hopped clones revealed integrations throughout the respective LADs (114/124 and 61/89), showing that LADs are not refractory to SB excision and re-integration. Instead, in these latter experiments, most integrations occurred within the LAD, with few hopping events to the neighbouring iLADs (10/124 and 28/89). We conclude that hopping of SB is efficient within both LADs and iLADs, but is rare between the two compartment types, presumably because these compartments are spatially separated.

*Efficient Cre recombination in selected clones.* From the thus obtained clones with *loxP* integrations throughout LAD1 or LAD2, we selected 12 for the generation of inversions and deletions. For each of these clones, we transiently expressed Cre, isolated 30-150 Cre-expressing subclones, and identified recombined subclones by PCR screening (see Methods). This revealed that the recombination efficiency was remarkably high, both for deletions and inversions, reaching up to 22% for *loxP* sites as much as 1.8 Mb apart (**Figure 1E**). Surprisingly, the recombination efficiency did not correlate with the distance between *loxP* sites (R = 0.22, p = 0.5).

### Only large deletions within LAD1 result in release from the NL

*Deletion series of LAD1: questions that we address.* We first focused on six deletions that consisted of increasingly large truncations of LAD1, combined with loss of a constant ∼630 kb part of the flanking iLAD (**Figure 2A**). Each of these deletions resulted in a new iLAD-LAD junction connecting an identical iLAD sequence with a different LAD1 sequence. This deletion series allowed us to address two questions. First, how are NL interactions of sequences near the new junctions altered compared to the wild-type context? Second, is full integrity of LAD1 required for its NL interactions, and if not, which parts are needed?

**Figure 2:**
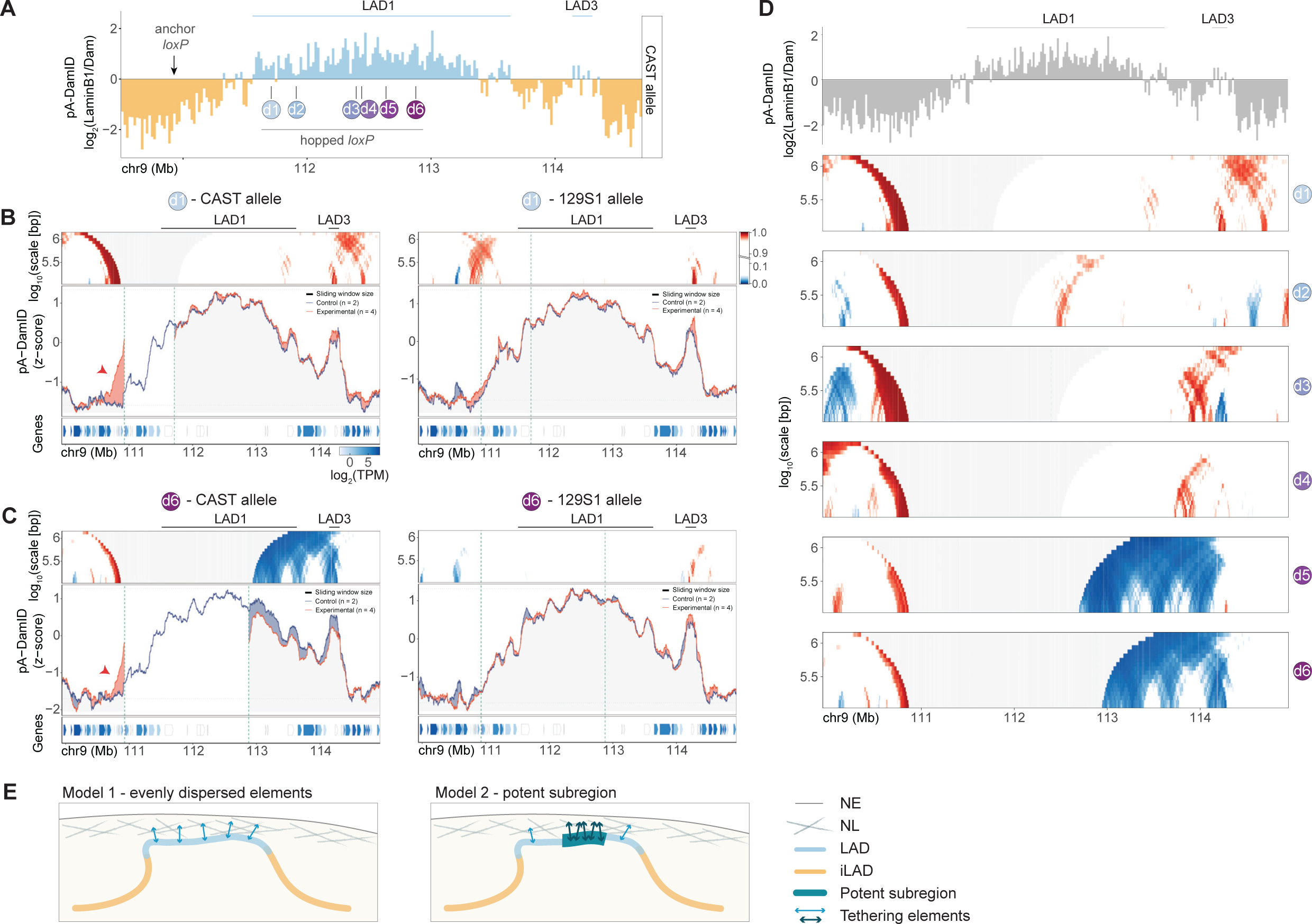
Large LAD1 deletions only result in LAD release from the NL. **(A)** CAST allele-specific pA-DamID track of NL interactions (bin size 20 kb), together with the positions of the six *loxP* integrations within LAD1 that were used in combination with the anchor *loxP* to generate deletions d1 through d6. **(B-C)** Changes in NL interactions resulting from deletions. Each of the four panels consists of three sub-panels. Bottom sub-panel: gene annotation track as in Figure 1B. Middle sub-panel: pA-DamID tracks (z-score) of NL interactions in non-recombined (“Control”, blue line) and recombined cells (“Experimental”, red line). n indicates number of independent biological replicates that were combined. Noise was suppressed by a running mean filter of indicated window size. Shading between the lines corresponds to the colour of the sample with the highest value. Horizontal dashed lines mark the 5^th^ and 95^th^ percentiles of genome-wide pA-DamID values. Vertical dashed lines mark the coordinates of the *loxP* recombination sites. For the experimental dataset, background noise coming from the deleted region on the CAST allele was filtered out. Top sub-panel: domainogram (Brueckner et al., 2020); for every window of indicated size (vertical axis) and centred on a genomic position (horizontal axis), the pixel shade indicates the ranking of the change in pA-DamID score (Experimental minus Control) in this window compared to the genome-wide changes in pA-DamID scores across all possible windows of the same size. Blue: pA-DamID score is highest in control samples; red: pA-DamID score is highest in experimental samples; grey: no data, the change in pA-DamID score cannot be calculated over the deleted region. Data are shown separately for the CAST (left panels) and 129S1 (right panels) alleles. (B) Smallest deletion d1; (C) largest deletion d6. Arrowheads mark an iLAD region that gains NL interactions in recombined cells. **(D)** Comparison of CAST-allele-specific domainograms of all six deletions involving LAD1 (bottom six panels). Top panel shows same pA-DamID track as in (A) for reference. **(E)** Two possible models for LAD1 interactions with the NL. Left: NL-tethering elements are evenly distributed throughout the LAD; right: critical subregions have stronger NL-tethering activity. NE: nuclear envelope.

*How data are generated and plotted.* For each deletion we selected two independent recombined clones and generated detailed maps of NL interactions by pA-DamID ^45^. The parental clone, (with hopped but non-recombined *loxP*) was used as control. Inspection of the allele-specific pA-DamID read counts around the recombination sites confirmed the expected deletions in the CAST allele but not in the 129 allele of all recombined clones (**Figure S2**). For each deletion, we visualized effects of the deletions on NL interactions in two ways: a plot of the pA-DamID tracks of control (blue) and recombined (red) clones (**Figure 2B-C**, middle panels); and a multi-scale “domainogram” ^41^. that highlights regions with increases (red) or decreases (blue) in NL contacts that are unlikely to be due to random noise (**Figure 2B-C**, top panels; see legend for detailed explanation). The independent replicate clones yielded similar NL association patterns for all recombinations (**Figure S3A**), therefore we combined the data of both clones.

*Local gain of NL interactions of iLAD sequences near the junction.* The smallest deletion that we generated was 784 kb long and included ∼150 kb of LAD1 (recombination d1). Strikingly, near the newly created junction, this resulted in increased NL interactions of the formerly iLAD sequence over a range of ∼190 kb (**Figure 2B, left panel, arrowhead**). We did not observe similarly strong changes for the 129S1 allele in the same cells (**Figure 2B, right panel**), demonstrating that the changes in NL contacts on the CAST allele are directly caused by the deletion. These data indicate that NL interactions spread *in cis* across the junction. In contrast, the deletion did not detectably affect the NL contacts of the intact part of LAD1 (**Figure 2B, left panel**). From this we conclude that the deleted sequences are not required for the formation of LAD1.

*Same phenotype for LAD1 deletions up to ∼880kb.* Next, we analysed recombination d2, d3 and d4, in which stretches of about 350 kb, 840 kb and 880 kb of LAD1 were deleted, respectively. Despite these much larger deletions, we observed a similar phenotype as described for recombination d1 (**Figure 2D; Figure S3B-C-D**). In each case, the iLAD region near the new junction gained NL interactions. Importantly, also in these three deletions, the NL interactions of the remaining parts of LAD1 remained unaffected. Thus, ∼40% of LAD1 length can be deleted without detectable impact on the association of the remaining ∼60% with the NL, showing that LAD1 is very robust.

*Larger deletions partially destabilise LAD1*. In contrast, deletions causing even larger truncations of LAD1 (about 1.1 Mb and 1.3 Mb for recombination d5 and d6, respectively) led to more substantial changes in the NL interaction pattern. While the iLAD region near the new junctions still gained NL interactions – as we observed for the smaller deletions – the remaining parts of LAD1 now showed significantly reduced NL interactions (**Figure 2C-D; Figure S3E**). Nevertheless, the NL interactions of LAD1 were never completely lost. Interestingly, in recombination d6, the NL interactions of LAD3 were also somewhat reduced, suggesting crosstalk between these LADs.

*Summary of key results.* Direct comparison of the domainograms of the six deletions (**Figure 2D**) provides a summary of the main changes in NL interactions. All six deletions caused spreading of NL interactions into the former iLAD region near the junction, over a range of ∼190-260 kb. This effect was most prominent for recombination d3, possibly because the breakpoint of this deletion is located in the part of LAD1 that has the strongest NL interactions. Furthermore, LAD1 is remarkably robust. Deletion of nearly half of LAD1 does not impair its association with the NL (recombination d1 to d4). However, larger deletions resulted in a significant – albeit incomplete – release of the remaining LAD sequences from the NL (recombination d5 and d6).

*Two models for LAD formatio*n. These results may be explained by two models. In the first model, NL interactions are mediated by a multitude of elements (which may be DNA sequence elements or a chromatin modification) distributed evenly throughout each LAD. Individually, these elements may interact only weakly with the NL, but collectively they result in high avidity (**Figure 2E, left**). Only when the total mass of the LAD is reduced below a critical threshold the NL interactions may become less robust, as we observed for LAD1 in recombination d5 and d6. In a second model, tethering elements are dispersed unevenly throughout LADs, with critical subregions harbouring either a higher density of elements or elements with a stronger NL tethering activity (**Figure 2E, right**). Because only recombination d5 and d6 lead to reduced NL interactions, the region located between deletion breakpoints 4 and 5 (R4-5) may constitute such a critical subregion in LAD1. We note that the remainder of LAD1 in recombination d5 and d6 would still harbour at least some tethering elements, since its contacts with the NL are not entirely lost.

### Inversion of LAD1 points to multiple tethering elements

*Breaking LAD1 into two halves by inversion.* To test the proposed models for LAD formation, we used our *loxP*-SB system to create an inversion that splits LAD1 into two halves of roughly equal size (recombination i7). If the NL contacts are driven by multiple elements uniformly distributed throughout the LAD, the two halves should both maintain interactions with the NL, although possibly with lower avidity. This inversion splits LAD1 into two parts (LAD1.1 and LAD1.2) that are separated by ∼630 kb of iLAD sequences (**Figure 3A**; **Figure 3B, top panel**). Again, we selected two recombined clones (**Figure S4A**) and performed pA-DamID. To facilitate the interpretation of the NL interaction pattern after recombination, we plotted the pA-DamID tracks and associated domainograms using chromosomal coordinates of the inverted genome, and re-plotted the non-recombined control data to this inverted genome. The results show that both LAD1 halves remained associated with the NL after the inversion. However, the inverted LAD1.1 part exhibited weakened NL interactions, particularly near the new iLAD junction (**Figure 3B, dashed arrow; Figure S4B**), while the non-inverted LAD1.2 part remained associated with the NL (**Figure 3B, solid arrow**). These results indicate that both LAD1 halves contain tethering elements, which differ in strength or quantity.

**Figure 3.**
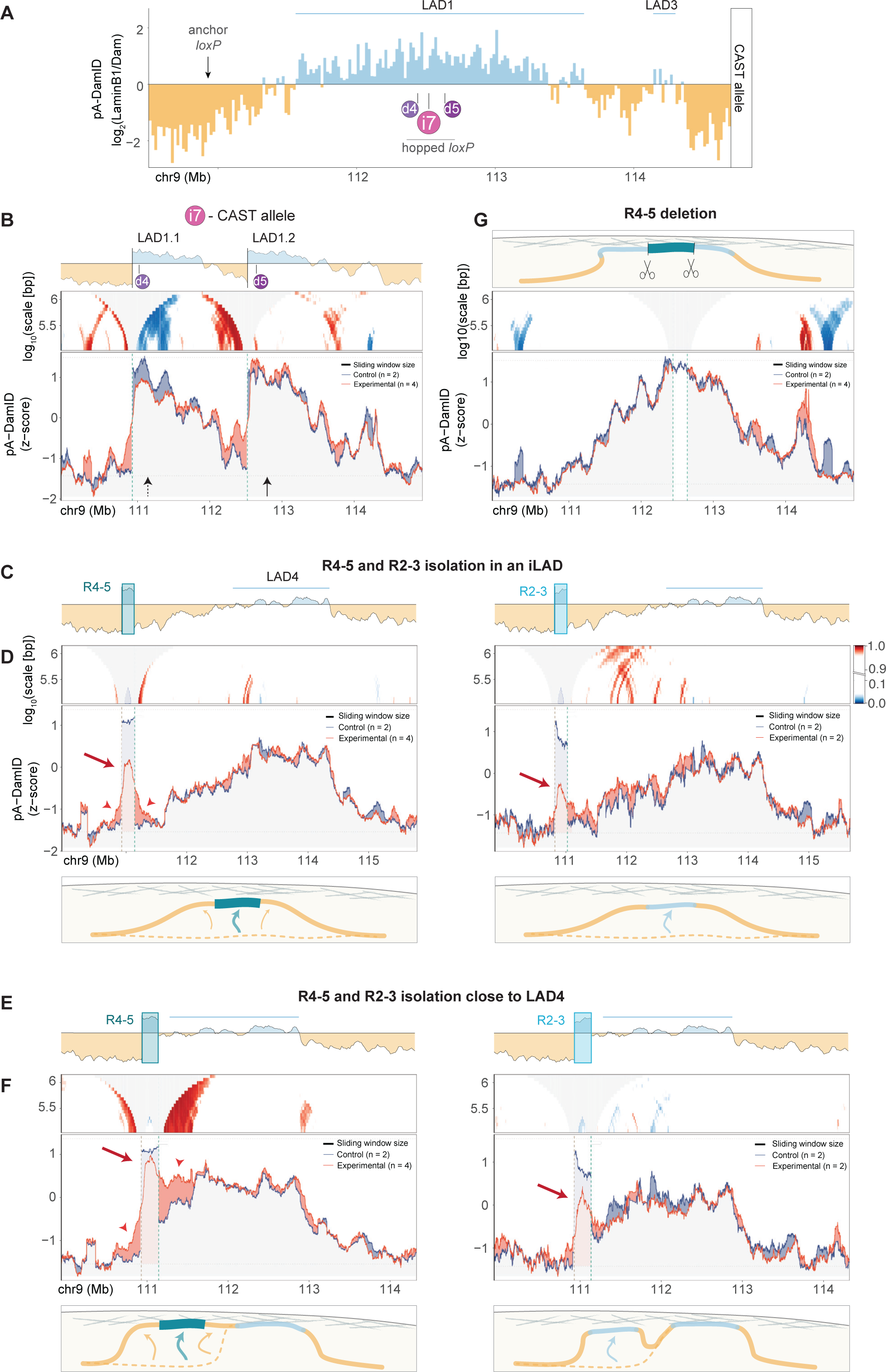
NL-tethering subregions in LAD1 are autonomous but not essential. **(A)** CAST allele-specific pA-DamID track of NL interactions in WT cells, together with the positions of three *loxP* integrations within LAD1 that were used in combination with the anchor *loxP* to generate rearrangements d4, d5 and i7. **(B)** Effect of splitting LAD1 into two halves. Top panel: cartoon of a pA-DamID track of NL interactions of the CAST allele, to illustrate the i7 inversion. Middle and bottom panels: domainogram (middle) and pA-DamID tracks (bottom) of CAST allele NL interactions comparing non-recombined and recombined cells from recombination i7. Data are plotted using the chromosomal coordinates of the inverted genome. Domainogram colour-key is displayed in (D). Dashed arrow indicates LAD1.1, which shows reduced NL contacts upon recombination; solid arrow indicates LAD1.2, whose NL interactions are largely unaffected by the recombination. **(C-D)** Effects of isolating R4-5 (left) and R2-3 (right) in an iLAD. Rearrangements were generated by sequential inversion and deletion, see Figure S5. (C) Cartoon of pA-DamID tracks of NL interactions of the CAST allele to illustrate the rearrangements. (D) Same plots as in (B) for the CAST allele of cells with hopped but not recombined loxP (“Control”, blue) and rearranged cells (“Experimental”, red) in which R4-5 and R2-3 regions have been isolated within an iLAD. pA-DamID tracks and associated domainograms are plotted using chromosomal coordinates of the rearranged genome. Arrows indicate NL interactions of R4-5 and R2-3 upon their isolation, arrowheads mark R4-5 neighbouring sequences gaining NL interactions. Bottom panels: schematics of NL interaction changes upon R4-5 and R2-3 isolation in an iLAD. For legend, see Figure 2E. **(E-F)** Same as (C-D) but after insertion of R4-5 and R2-3 close to LAD4. **(G)** No detectable effect of deletion of R4-5 from LAD1. Top panel: schematic of the CRISPR/Cas9-mediated deletion of R4-5. Bottom panels: Same plots as in B for the CAST allele of WT and CRISPR/Cas9-edited cells (“Experimental”, red) in which R4-5 has been deleted.

*Possible tethering role for a sub-segment of R4-5*. Interestingly, this inversion splits R4-5 into two parts of 80 and 120 kb that were divided over LAD1.1 and LAD1.2, respectively (**Figure 3A-B**). Because the 120 kb segment remained associated with the NL while the 80 kb segment showed decreased NL interactions after the inversion, it is likely that the 120 kb segment (positioned between breakpoints 7 and 5) harbours most of the tethering activity of R4-5.

*Spreading into neighbouring iLAD sequence*. At the two new LAD-iLAD junctions created by this inversion, we observed again increased NL interactions of the former iLAD sequences (**Figure 3B**). This effect was most pronounced for the iLAD region wedged between LAD1.1 and LAD1.2. Thus, as with the deletions described above (**Figure 2**), some spreading of NL interactions occurs across the new LAD-iLAD junctions.

### LAD1 subregion R4-5 can drive NL interactions

*R4-5 inserted in an iLAD* environment *has partially autonomous NL affinity.* To further test the capacity of R4-5 as a tethering element, we devised a strategy to isolate it from its native LAD context and relocate it into an iLAD environment. We then tested whether it could autonomously interact with the NL. We created such a rearrangement in two steps (**Figure S5A**). First, we used our *loxP*-SB system to generate an inversion within LAD1 that moved R4-5 directly next to iLAD sequences. Next, we used CRISPR/Cas9 editing ^46-48^ to delete a 3.7 Mb segment that included all LAD1 sequences except R4-5, resulting in the positioning of R4-5 between two iLAD regions of about 1.2 Mb and 1.6 Mb (**Figure 3C, left**). For comparison, we chose another LAD1 subregion, located between breakpoints 2 and 3 (named R2-3), of the same length as R4-5 and inserted in the same location by a similar approach (**Figure 3C, right; Figure S5B**). Again, we selected clones for each rearrangement (**Figure S6A-B**), mapped their NL interactions by pA-DamID and compared them to wild-type cells. The results show that R4-5 has a modest but significant ability to autonomously interact with the NL, compared to the flanking iLAD regions (**Figure 3D, left; Figure S6C-D;** p < 2.22e-16, Wilcoxon text). Interestingly, R2-3 also interacts with the NL (p = 5.3e-4), but it does so less frequently than R4-5 (p = 3.2e-7) (**Figure 3D, right, Figure S6D**). The flanking iLAD sequences gained NL interactions when close to R4-5 only, although modestly.

*Cooperativity with a neighbouring LAD.* Although these results point to some degree of autonomous NL tethering by R4-5, its NL interactions in the iLAD context were weaker than those in the original LAD1 context (**Figure 3D**). Possibly, R4-5 requires cooperative interactions with other LAD elements for more efficient NL tethering. To test this, we isolated R4-5 and R2-3 regions in close vicinity of LAD4 (**Figure 3E; Figure S5; Figure S6E**). Interestingly, in this context, R4-5 exhibited nearly as much interaction with the NL as in its wild-type LAD1 context (**Figure 3F, left panel; Figure S6F**). Moreover, R4-5 enhanced the NL association of flanking LAD4 sequences over about 400 kb (**Figure 3F, left panel**). The iLAD sequences near the other junction also gained NL interactions, comparable to what we previously observed in the deletion series (recombination d2, d5, d6) (**Figure 2D**). In contrast, moving R2-3 close to LAD4 only slightly strengthened its NL interactions compared to the iLAD context, and it did not detectably boost NL contacts of its flanking sequences (**Figure 3F, right panel**). Thus, R4-5 and R2-3 may differ not only quantitatively in their autonomous ability to interact with the NL, but also in their ability to boost NL contacts of flanking LAD regions.

### R4-5 is not essential for NL interactions of entire LAD1

*Deletion of R4-5 has no effect on LAD1.* Next, we tested how much R4-5 contributes to NL interactions of LAD1, where it is normally located. For this we precisely excised R4-5 from LAD1 by means of CRISPR/Cas9 editing with guide RNAs specific for the CAST allele. pA-DamID data confirmed the expected deletion in the CAST allele in two independent clones (**Figure S6G**). Surprisingly, R4-5 deletion had no impact on LAD1 association with the NL (**Figure 3G; Figure S6H**). Thus, in the context of the entire LAD1, R4-5 is not essential, underscoring that NL interactions of LAD1 are redundantly controlled by multiple elements.

### LAD2 also harbours a boosting region

*Splitting of LAD2 points to a boosting region*. We next investigated whether the mechanisms driving LAD1 formation are also involved in the formation of the smaller LAD2. To test this, we generated an inversion with a breakpoint within LAD2, inverting one third of its sequence (recombination i10) (**Figure 4A**). As a result, LAD2 is split in two, with the inverted part being moved close to LAD1, about 1.2Mb away from its original location. We will refer to the non-inverted part as LAD2.1 and to the inverted part as LAD2.2 (**Figure 4B, top panel**). We performed pA-DamID on two recombined clones (**Figure S7A**) as well as on the parental one. Upon recombination, LAD2.2 showed slightly weakened NL interactions, but at the same time it promoted NL association of its new neighbouring iLAD sequences (**Figure 4B, solid arrow; Figure S7B**). LAD2.1 lost NL interactions more substantially (**Figure 4B, dashed arrows**), but it still contacted the NL to some extent. The observations that LAD2.2 triggers NL association of its neighbouring sequences and that the NL interactions of LAD2.1 are weakened upon inversion of LAD2.2, points to the identification of LAD2.2 as another NL interaction boosting region. Furthermore, both subregions must harbour NL tethering elements, as they do not completely lose their NL contacts after the inversion. Due to the high repetitiveness of LAD2 sequences, we could not perform any CRISPR/Cas9 deletion for further detailed dissection.

**Figure 4.**
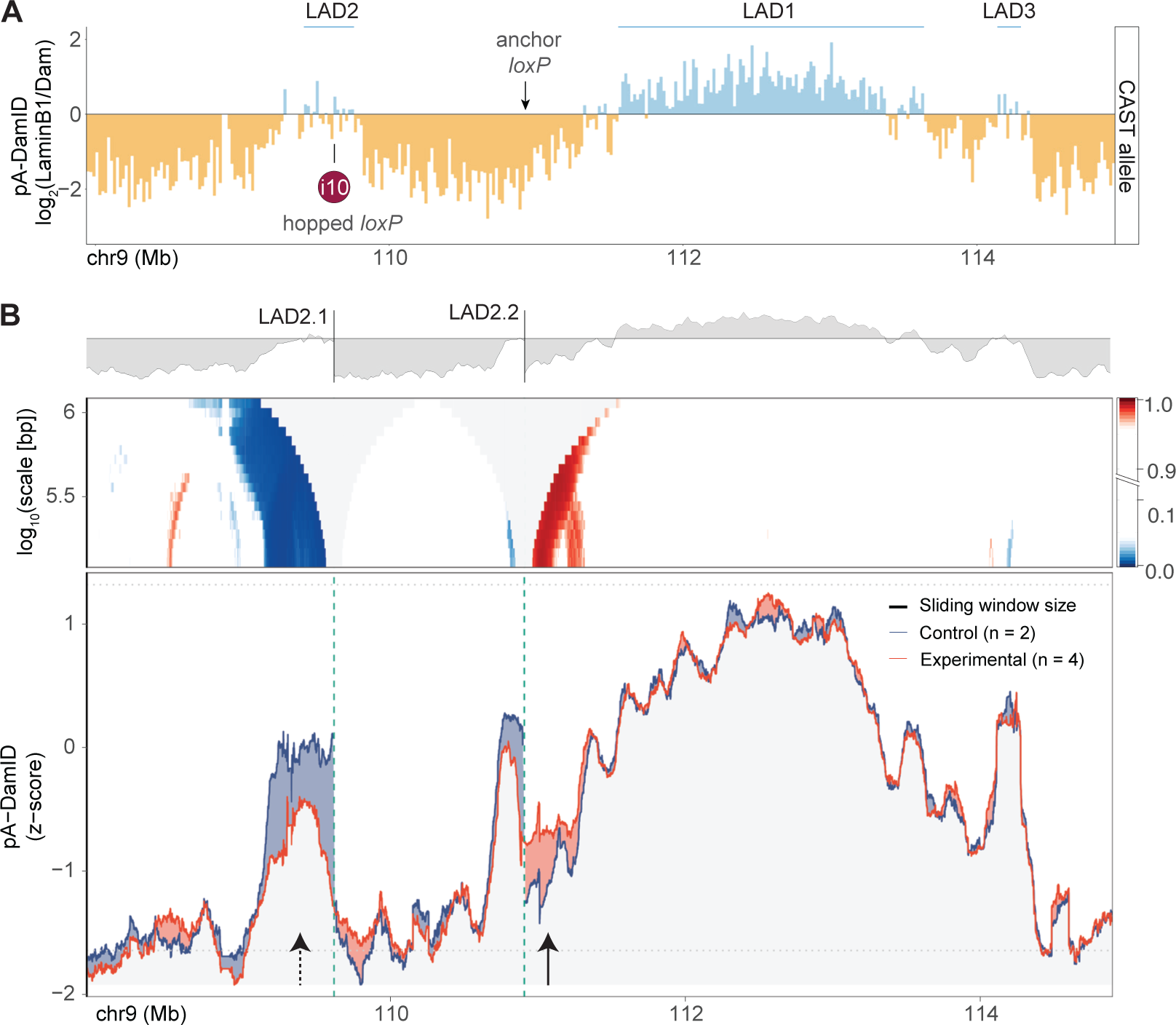
LAD2 contains a subregion that boosts NL interactions. **(A)** CAST allele-specific pA-DamID track of NL interactions in WT cells, together with the position of one *loxP* integration within LAD2 that were used in combination with the anchor *loxP* to generate rearrangements i10. **(B)** Top panel: cartoon of a pA-DamID track of NL interactions of the CAST allele, to illustrate the i10 inversion, which divides LAD2 into two segments that are separated by a large iLAD region. Middle and bottom panels: domainogram (middle) and pA-DamID tracks (bottom) of CAST allele NL interactions comparing non-recombined cells (“Control”) to cells with inversion i10 (“Experimental”). Data are plotted using the chromosomal coordinates of the inverted genome. Dashed arrow indicates LAD2.1, which lose NL contacts upon recombination; solid arrow indicates LAD2.2-flanking iLAD sequences, which gain NL contacts upon recombination.

### No obvious sequence features that explain boosting regions

*No enrichment of cKrox and YY1 motifs.* We examined the boosting regions R4-5 and LAD2.2 for sequence features that might explain their affinity for the NL. Earlier studies suggested that cKrox and YY1 transcription factors participate in the tethering of ectopic LAD fragments to the NL ^26,28^. However, we found that neither the motif originally assigned to cKrox (erroneously inferred from the *Drosophila* homolog Trl) ^26^, nor the up-to-date cKrox binding motif ^49^, nor the YY1 motif ^49^ are enriched in R4-5 and LAD2.2 (**Figure S8A**). Genome-wide, the cKrox and YY1 motifs are even depleted in LADs compared to iLADs and their density in the two boosting regions is similar to the median value across all LADs (**Figure S8B**). It is thus unlikely that these motifs mediate the NL attachment of R4-5 and LAD2.2.

*No shared repeat enrichment or elevated AT-content.* We also considered a wide range of repeat elements, some of which are known to be enriched in LADs genome-wide ^6,50,51^. While LAD2.2 has somewhat higher densities of several types of repeats compared to LAD2.1, this is not the case for R4-5 compared to the remainder of LAD1 (**Figure S8C-D**). Finally, we analyzed overall AT-content, which is known to be high in LADs ^9^. Within LAD1, R4-5 has indeed a relatively high A/T-content compared to the remainder of LAD1, but this enrichment of A/T-rich sequences is not observed in LAD2.2 (**Figure S8E**). We conclude that there is no obvious shared sequence characteristic that stands out in the two boosting regions.

### Synergy between LAD1 and LAD2 requires linear proximity

*Do neighbouring LADs promote each other’s NL interactions?* LADs are known to interact with each other within the B compartment as defined by Hi-C mapping ^52-54^ and can be engaged in physical ‘cliques’ ^32^. Furthermore, single-cell DamID mapping has indicated that neighbouring LADs tend to interact with the NL in a coordinated fashion ^23^. We thus wondered whether neighbouring LADs could promote each other’s interaction with the NL.

*Deletion of LAD2 does not alter NL contacts of LAD1.* We first tested whether LAD2 participates in the association of LAD1 with the NL. We again used our *loxP*-SB system to delete the entire LAD2 from the CAST allele of mESCs (recombination d11) (**Figure 5A**). We selected recombined clones (**Figure S9A**) and performed pA-DamID. This revealed that the interactions of LAD1 with the NL are not affected by LAD2 deletion (**Figure 5B; Figure S9B**).

**Figure 5.**
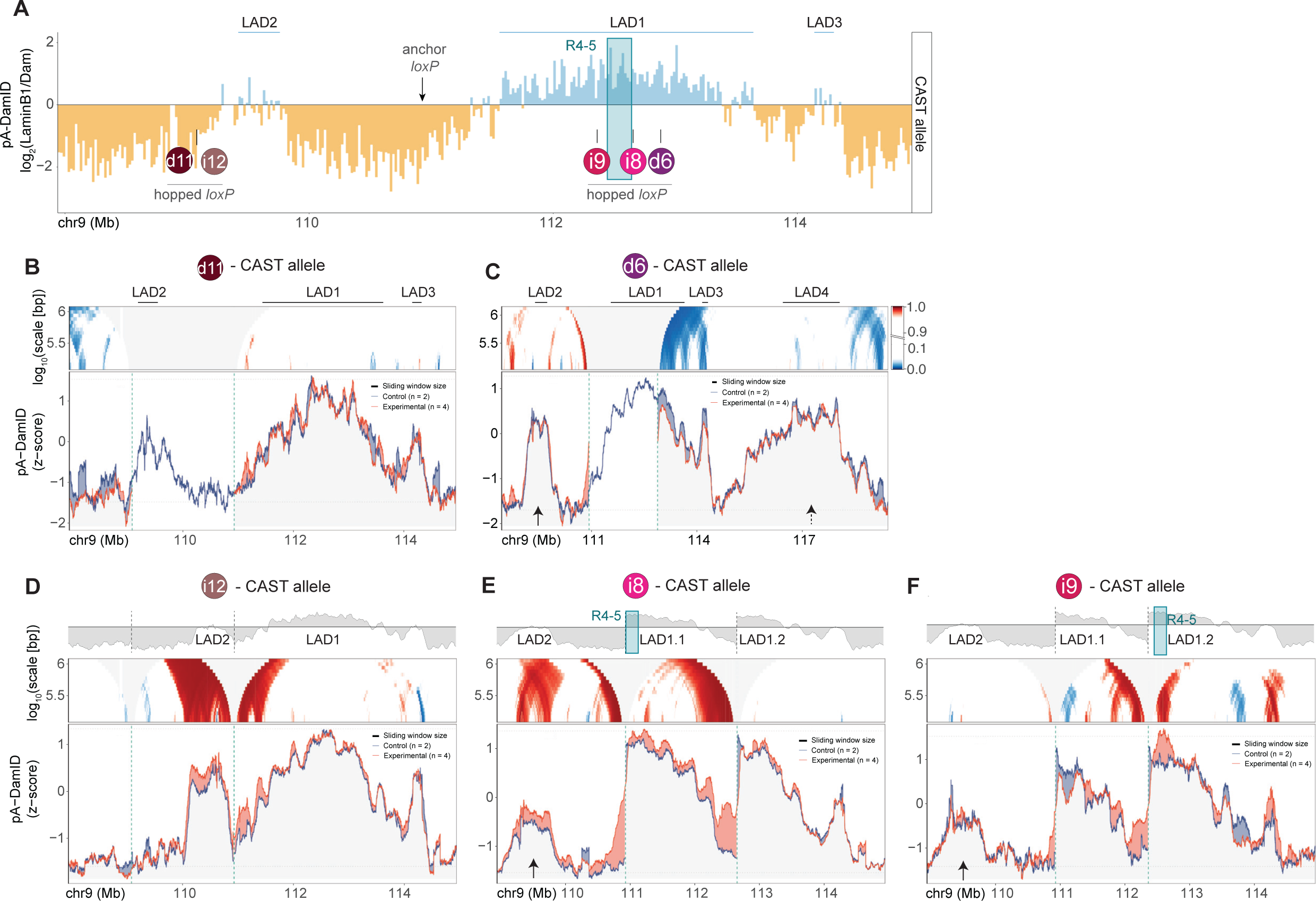
Synergy between LAD1 and LAD2 when in close proximity. **(A)** CAST allele-specific pA-DamID track of NL interactions in WT cells, with the positions of five *loxP* integrations that were used to generate genomic deletions (d6, d11) or inversions (i8, i9, i12). **(B)** Deletion of LAD2 does not affect LAD1. Domainogram and pA-DamD tracks for the CAST allele of non-recombined and recombined cells from recombination d11. **(C)** Deletion of a large part of LAD1 does not affect LAD2 and LAD4. Same plot as in Figure 2C, left, but for a wider window. Solid and dashed arrows indicate LAD2 and LAD4 respectively. **(D-F)** Top panels: cartoon of a pA-DamID tracks of NL interactions, to illustrate inversions i12 (D), i8 (E) and i9 (F). Middle and bottom panels: domainograms (middle) and pA-DamID tracks (bottom) comparing the CAST allele of non-recombined (“Control”) and respective recombinations (“Experimental”) as indicated. In (E) and (F), arrows indicate LAD2. Domainogram colour-key in (C) also applies to (B, D-F).

*LAD1 does not help LAD2.* Because LAD1 is longer and stronger than LAD2, it is more likely that it participates in NL contacts of LAD2. To test this, we revisited recombination d6, the largest deletion we generated in LAD1 (**Figure 2C**). Although the remainder of LAD1 is much smaller and weaker, LAD2 remained mostly unaffected by this LAD1 deletion (**Figure 5C, solid arrow; Figure S9C**). We could not detect any change in LAD4 either (**Figure 5C, dashed arrow**). Thus, these experiments suggest that LAD1 and LAD2 independently interact with the NL.

*Synergy between LADs depend on their distance.* We wondered whether the 2 Mb long, transcriptionally highly active iLAD region between LAD1 and LAD2 could prevent crosstalk between the two LADs. To shorten the LAD1-LAD2 linear distance, we created an inversion that brought LAD2 within ∼700 kb from LAD1 (**Figure 5A, recombination i12; Figure 5D, top panel; Figure S9D**). Strikingly, upon this inversion, both LAD2 itself and the iLAD sequences wedged between LAD1 and LAD2 gained NL interactions (**Figure 5D; Figure S9E**). LAD1 did not gain NL interactions, though. These results suggest that, when LAD1 and LAD2 are close enough, LAD2 benefits from LAD1’s proximity and interacts more strongly with the NL.

*R4-5 can promote LAD1-LAD2 crosstalk*. To determine which component of LAD1 was necessary for enhancing the NL interactions of LAD2, we took advantage of recombination i8 and i9 in which LAD1 is split in two (**Figure 5A; Figure S5**). In recombination i8, R4-5 is the closest to LAD2 in the linear space, while in recombination i9 this is the case for R2-3. We performed pA-DamID on the recombined and parental clones (**Figure S6A**). We will refer to the inverted and non-inverted halves of LAD1 as LAD1.1 and LAD1.2 respectively. When R4-5 was positioned close to LAD2 in recombination i8, LAD2 gained lamina interaction (**Figure 5E; Figure S9F**). However, LAD2 did not gain lamina interaction in recombination i9, when R2-3 was repositioned in its proximity (**Figure 5F; Figure S9G**). We note that recombination i9 shows several other gains and losses of NL interactions that cannot be easily explained; apparently the inversion causes multiple changes in LAD1.2 and nearby regions. Together, these data suggest that the boosting region R4-5 has the ability to promote crosstalk between LADs. We cannot rule out that LAD length plays a role in the crosstalk, because LAD1.1 is longer in recombination i8 than in recombination i9.

*LAD1.1-LAD1.2 crosstalk*. Another interesting feature of recombination i8 is that the NL interactions of LAD1.2 remained unaffected even though it is no longer contiguous with LAD1.1, but separated from it by ∼0.5 Mb of iLAD sequence (**Figure 5E**). This contrasts with recombination d5, in which the remainder of LAD1 (which corresponds to LAD1.2 in recombination i8) showed reduced NL interactions when LAD1.1 was deleted. This result suggests that LAD1.1 can enhance NL interactions of LAD1.2 even when they are separated by ∼0.5 Mb of iLAD sequences.

*Conclusion*. In conclusion, these data indicate that synergy can occur between two neighbouring LADs, provided that they are sufficiently close to one another in the linear space.

### Changes in NL interactions are partially mirrored by changes in H3K9me3

*pA-DamID of H3K9me3.* LADs often coincide with large domains of repressive histone marks such as H3K9me3 ^16,23,27,55,56^. We therefore investigated whether the observed changes in NL interactions were mirrored by changes in H3K9me3, by performing pA-DamID of this histone mark in several recombined and control cells. In wild-type cells, H3K9me3 indeed correlates substantially, although not perfectly, with NL interactions (**Figure S10A**, genome-wide Pearson correlation R = 0.61). Likewise, LAD1 is enriched in H3K9me3, although the fine-scale pattern differs at some positions from that of NL interactions (**Figure S10B**).

*NL interaction spreading can be accompanied by H3K9me3 spreading.* We first focused on recombination d3, particularly on the iLAD region that showed increased NL interactions when fused to part of LAD1 (**Figure 2D**). In recombined cells, this region also gained H3K9me3 (**Figure 6A; Figure S10C**). This increase in H3K9me3 extended over ∼200 kb but was more modest than the NL interaction gain, perhaps because these iLAD sequences already had substantial H3K9me3 levels in non-recombined cells. In recombination d6, in which the spreading of NL interactions across the new iLAD-LAD junction was less pronounced (**Figure 2C-D**), the increase in H3K9me3 was more subtle than in recombination d3 (**Figure 6B; Figure S10D**). Stronger local increases of H3K9me3 were detected at the iLAD-LAD junction in recombination i7 (**Figure 6C; Figure S10E**). Thus, at least in this region, local spreading of NL interactions and H3K9me3 appeared to correlate at the iLAD-LAD junction.

**Figure 6:**
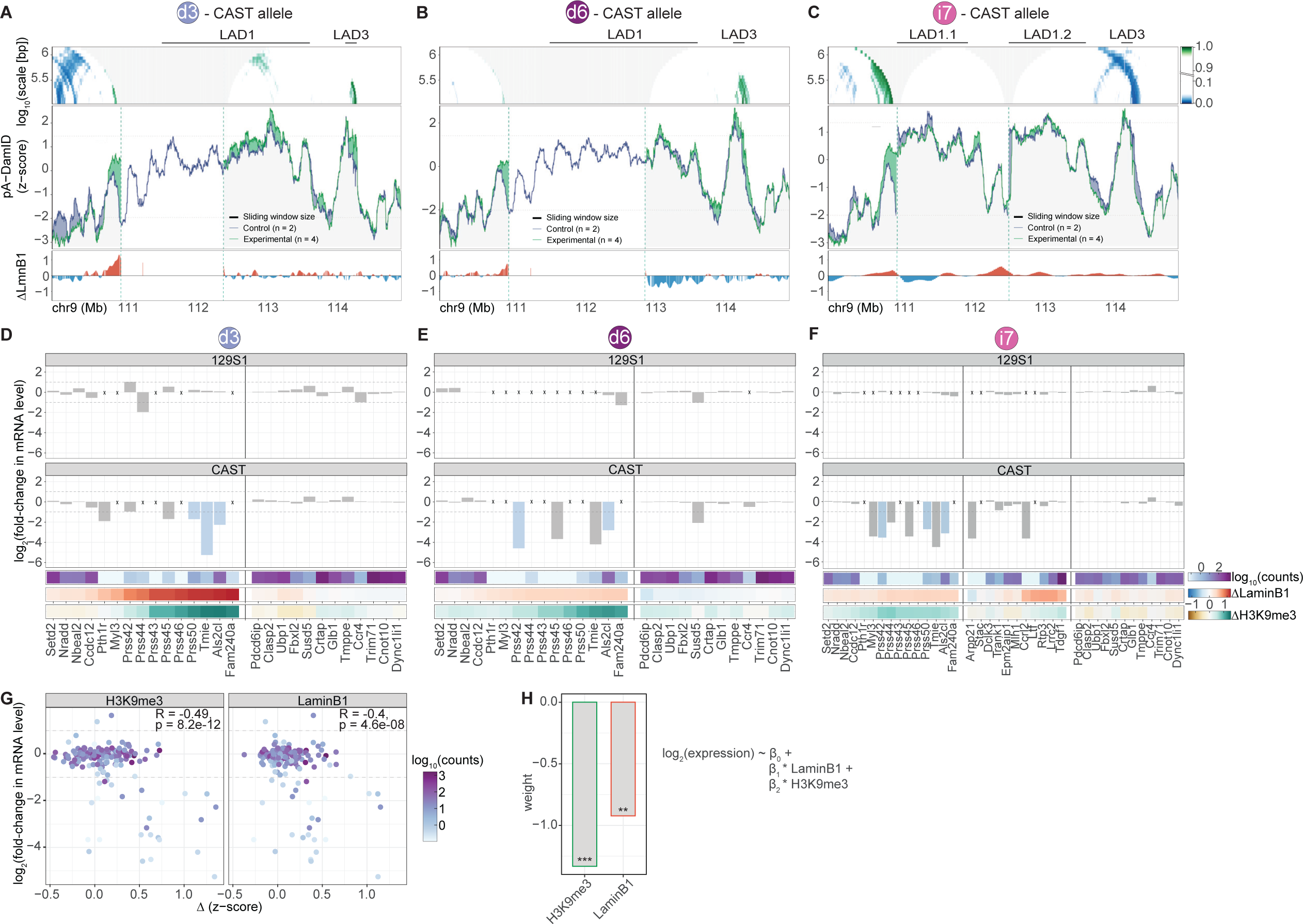
Spreading of NL contacts is partially mirrored by H3K9me3 and often linked to gene repression. **(A-C)** Changes in H3K9me3 caused by recombinations. Top and middle panels: domainograms and pA-DamID tracks showing CAST allele-specific H3K9me3 levels in control (blue) and recombined cells (“experimental”, green) for recombination d3 (A), d6 (B), and i7 (C). Bottom panels: tracks showing the change in NL interactions in the recombined clone, compared to the control (ΔLmnB1, recombined minus control). Data were smoothed. Dashed vertical lines mark positions of the recombined *loxP* sites. In panel (C) data are plotted using coordinates of the recombined genome. **(D-F)** Changes in gene expression in recombinations d3, d6, and i7. Bargraphs show allele-specific log_2_ fold-change in gene expression between the control and one recombined clone from recombination d3 (D), d6 (E) and i7 (F), for each gene around the recombined sequences. Genes are ordered by their chromosomal coordinates. Vertical lines represent the location of *loxP* recombination sites. Significant differentially expressed genes are coloured in blue (p-value corrected for multiple testing < 0.05 and fold-change > 2). Horizontal dashed lines mark log_2_(fold-change) of 1 and -1. Crosses: no detectable mRNA in the control cell line (with hopped but non-recombined *loxP*). Heatmaps underneath the bargraphs show for each gene (from top to bottom): (i) expression level in WT F1 hybrid mESCs, (ii) change in NL interaction pA-DamID score after recombination (recombined minus control), (iii) change in H3K9me3 pA-DamID score after recombination (recombined minus control). **(G)** Correlation between changes in gene expression and the change in H3K9me3 levels (left) or NL interactions (right). Aggregate plot shows the combined data of the CAST alleles of all genes depicted in D, E, F and Figure S10G. Each dot represents a gene in one tested recombination. Dots are coloured based on the gene expression level in WT F1 hybrid mESCs. Large dots represent significant differentially expressed genes. **(H)** Linear model of changes in gene expression as function of changes in both NL interactions and H3K9me3. Barplot shows weights of the fitted linear model; asterisks: ***: p < 0.001, **: p<0.01.

*Partial uncoupling of H3K9me3 and NL interaction changes.* Despite these observed similarities, we also found instances where changes in NL interactions were not mirrored by H3K9me3 changes. For example, in recombination i7, the iLAD sequences wedged between LAD1.1 and LAD1.2 gained NL interactions (**Figure 3B**; **Figure 6C**) but not H3K9me3 levels (**Figure 6C; Figure S10E**). Furthermore, in recombination d6, the reduction in NL interactions of the remaining part of LAD1 (**Figure 2C**) was not detectably accompanied by a lowering of H3K9me3 levels (**Figure 6B**). Likewise, in the recombined clones with partial or complete LAD2 inversion (i10 and i12 respectively) no significant changes in H3K9me3 levels were detected in regions with losses or gains of NL interactions (**Figure S10F-G**). We conclude that changes in NL interaction levels caused by chromosomal structural variants are only partially mirrored by changes in H3K9me3.

### Spreading of NL contacts and H3K9me3 is accompanied by transcriptional repression

*mRNA-seq analysis of recombination d3.* Many studies have found that the localisation of genes at the NL is correlated with a lack of transcriptional activity ^6,7,15-19^, but so far only a few reports have pointed to a causal role of NL proximity in mammalian cells ^18,19^. We therefore examined whether the observed changes in NL interactions, H3K9me3 levels, or both, were accompanied by changes in gene expression as detected by mRNA sequencing. We first focused on recombination d3. We considered every gene within the window plotted in **Figure 6A**, and defined differentially expressed genes as those showing a statistically significant change in mRNA abundance as well as a fold-change of more than 2.

*Gain of NL interactions and H3K9me3 is linked to gene repression.* Outside from the deleted region, we detected three significantly downregulated genes (*Prss50, Tmie* and *Als2cl*) on the recombined chromosome (**Figure 6D**). Strikingly, these genes are close to the LAD1 junction and showed the strongest gain in both NL association and H3K9me3 levels after recombination (**Figure 6D, bottom panels**). Only one gene (*Fam240a*) is closer to the junction, but this gene is silent in wild-type cells and hence cannot be further downregulated. No significantly differentially expressed genes were detected on the 129S1 allele, demonstrating that the observed changes on the CAST allele were the direct consequence of the deletion *in cis* (**Figure 6D, top panel**).

*Weaker gain in NL interactions also lead to gene downregulation.* In recombination d6, gain in NL interactions and H3K9me3 levels were milder compared to recombination d3 (**Figure 6A-B-D-E**). Still, two genes were found significantly downregulated (*Prss42* and *Als2cl*) and two others showed substantial downregulation although not statistically significant (*Prss45* and *Tmie*) (**Figure 6E**). We conclude that a set of genes close to the deletion junction are responsive to the NL environment and are downregulated upon gain of NL interactions and H3K9me3.

*No transcription activation of genes with reduced NL interactions.* In recombination d6 several genes inside the remainder of LAD1 showed reduced NL interactions (**Figure 6B-E**). However, we found no significant upregulation of these genes (**Figure 6E**). This could be because their association with the NL or their H3K9me3 levels are still too strong to be compatible with transcriptional activation. Alternatively, these genes could lack the potential for activation ^57^, for example because they require specific transcription factors that are missing in mESCs.

*Gene repression correlates more with H3K9me3 than NL interactions.* We extended this mRNA-seq to two recombined clones from recombination i7. The *Als2cl, Prss50 and Prss42* genes, which gained both NL interactions and H3K9me3 were significantly downregulated (**Figure 6F**). Several nearby genes followed the same trend, albeit non-significantly. However, none of the iLAD genes that gained NL interactions only (e.g. *Lrrc2*, *Rpt3*) were significantly downregulated. This suggested that H3K9me3 level is a more important determinant for gene repression than NL association. To analyse this systematically, we combined mRNA-seq data from above-mentioned deletions and inversions and additionally included inversions i10 and i12 (**Figure S10H**). We then plotted the fold-change in gene expression relative to the change in NL interactions or H3K9me3 levels (**Figure 6G**). The correlation was modestly stronger when expression fold-change was compared to H3K9me3 changes than to NL interactions changes (Pearson’s R = -0.49 and R = -0.4, respectively). A combined linear model indicated that changes in H3K9me3 are a slightly more important predictor of the changes in gene expression than changes in NL interactions (**Figure 6H**). However, the linear model only explains only about a quarter of the variance in gene expression (R^2^ = 0.27, p = 2.3e-13), suggesting that other unknown features also play a role. In conclusion, the inversions and deletions of iLAD/LAD sequences not only alter NL interactions, but also H3K9me3 and gene activity.

## DISCUSSION

*Summary of key results.* The very large sizes of LADs have made it difficult to unravel how they interact with the NL. In this study, we combined SB hopping and Cre-lox recombination to generate a large set of local deletions and inversions of LAD and iLAD sequences. Detailed mapping of NL interactions in the mutant clonal cell lines enabled us to uncover several principles of LAD - NL interactions. We show that LADs contact the NL via multiple LAD subregions that differ in NL tethering strength, in an intricate pattern of cooperativity and redundancies that underscore that LAD - NL interactions are multivalent. Potent subregions (here referred to as boosting regions) can autonomously interact with the NL and boost NL association of flanking sequences. Furthermore, we find that neighbouring LADs can synergistically interact with the NL, provided that they are close enough on the same chromosome.

*Necessity of the boosting region depends on LAD strength.* Of the two boosting regions that we identified, one is necessary for the formation of its corresponding LAD (LAD2) while the other one is not (LAD1). Indeed, when the LAD2 boosting region is moved away from its original location the remaining part of LAD2 loses NL interactions (**Figure 4**) while R4-5 is not necessary for the NL contacts of LAD1 (**Figure 3G**). Interestingly, in mESCs LAD1 has higher NL contact frequency compared to LAD2. We thus hypothesise that NL tethering of LAD1 involves several redundantly acting regions, while LAD2 mainly depends on a single region.

*Spreading of NL contacts implies active tethering*. At newly created LAD-iLAD junctions, we consistently observed spreading of NL interactions into the iLAD over 150-200 kb. This occurred both after we deleted the original LAD-iLAD border region (**Figure 2B-D**; **Figure 3B**) and after we inserted R4-5 into an iLAD (**Figure 3D**). NL interactions were even enhanced over ∼400 kb when R4-5 was inserted in an iLAD location close to a LAD (**Figure 3F**). An important implication of this spreading is that LAD-NL interactions are not a passive process, e.g., due to expulsion of transcriptionally silent chromatin from the transcriptionally active nuclear interior towards the NL. Rather, we propose that these interactions involve an active mechanism. As a consequence, neighbouring sequences are kept close to the NL as well.

*Spreading of NL interactions does not match persistence length.* We considered that the extent of this spreading could reflect the persistence length of the chromatin fibre: if the fibre is stiff over a long range, then any sequence that is linked to a NL-tethered sequence would also remain in close proximity to the NL over this range, which may be detected by pA-DamID. However, the persistence length of heterochromatin has been estimated to be only ∼100-200 nm ^58^, or ∼5-10 kb (assuming a compaction of 50 bp/nm ^59^). This is at least an order of magnitude shorter than the distance over which we observe spreading of NL contacts. An alternative explanation may be that NL contacts spread in a zipper-like manner. Evidence for such a mechanism was previously found in single-cell DamID maps, which indicated that NL interactions of LADs occur over stretches of several hundreds of kilobases ^23^. The molecular basis of such a zipper-like mechanism remains unknown, but it is consistent with our conclusion that LAD-NL interactions are multivalent.

*LAD spreading and new border formation.* A peculiar finding in the deletion series of LAD1 (**Figure 2**) is that the spreading into the flanking iLAD consistently extended over ∼150 kb, irrespective of the fused LAD segment. Remarkably, we detected spreading to a similar extend in recombination i7 (**Figure 3B**) and upon isolation of R4-5 in an iLAD environment (**Figure 3B**). A closer look in this iLAD region revealed that the spreading covered several lowly expressed genes and terminated around the first highly active gene. Previously, it was found that LAD borders are frequently demarcated by promoters of active genes ^6^, and that transcriptional activity can prevent NL interactions ^41,60^. We therefore propose that upon deletion of the original LAD1 border, the first highly expressed gene upstream of LAD1 acts as a new border that blocks further spreading of NL interactions.

*(i)LAD spreading depends on the flanking chromatin state*. LAD/iLAD border deletion may not always result in spreading of NL interactions. In CD4+CD8+ mouse thymocytes, deletion of a LAD border in the *Tcrb* locus induced the release of LAD sequences from the NL and activation of the displaced genes ^61^. In other words, LAD border deletion in the *Tcrb* locus resulted in iLAD spreading. The deleted sequence was directly flanked by an active enhancer marked by the active histone mark H3K27Ac and sequences that lost NL association concomitantly gained H3K27Ac ^61^. Another study found that activation of a long non-coding RNA gene enabled the detachment of a neighbouring gene from the NL ^62^. Thus, transcriptional activity can shape the pattern of NL interactions, and in some instances may limit the spreading of NL interactions.

*Synergy between neighbouring LADs.* Hi-C mapping studies revealed that LADs interact with each other within the B-compartment ^32,52-54^. Later on, polymer modelling suggested that LAD-LAD interactions are required for 3D genome folding ^10-14^. Yet, whether these homotypic interactions participate in the formation or stabilization of LADs themselves remained unclear. Our results indicate that LADs can cooperate to boost their association with the NL. Most likely this occurs through their physical interactions, which may effectively extend the multivalency of NL interactions beyond single LADs. Our data indicate that the efficacy of this mechanism is dependent on the linear distance between the LADs and on the presence of a boosting region like R4-5.

*Potential mechanism for NL affinity of boosting regions.* We identified two boosting regions, in LAD1 and LAD2. What could mediate their autonomous affinity for the NL? An obvious candidate would be a specific DNA sequence. Our results indicate that previously implicated sequence elements ^26,28^ are unlikely to play a role in mESCs, as they are not enriched in LADs genome-wide, nor strikingly abundant in the two boosting regions (**Figure S8**). This observation includes GAGA repeats, which were previously thought to interact with the NL via the DNA-binding factor cKrox ^26^. However, a role for cKrox could not be confirmed in another study ^27^, and this protein (now known as ZBTB17) was later found to bind a very different motif that is not enriched in LADs either. One of the two boosting regions (R4-5) has an unusually high AT-content. This fits with earlier findings that LADs (in particular constitutive LADs) are A/T-rich ^9^. Many proteins are known or predicted to bind to A/T rich sequences. Interactions of some of these proteins with A/T-rich boosting regions could mediate NL anchoring, but their identity remains to be established. However, the affinity of boosting regions for the NL could also be sequence-independent. These regions could adopt alternative non-B DNA conformations or specific supercoiling structures. Consistent with this hypothesis, Lamin B Receptor (LBR) mediates at least in part the peripheral positioning of heterochromatin ^63,64^ and was shown to preferentially interact with supercoiled DNA ^65^. Specific concentration of factors such as HP1 or modified histones could also mediate anchoring of boosting regions at the NL ^22,66^.

*Untangling NL association and gene repression.* Finally, we investigated the relationship between NL association, gene repression and deposition of the repressive histone mark H3K9me3. We show that NL association and H3K9me3 deposition are mediated by partially independent mechanisms, because changes in the former are only partially mirrored by changes in the latter. Moreover, gene expression changes correlate somewhat beher with H3K9me3 changes. This is reminiscent of what has been found in *Caenorhabditis elegans*, where transcriptional repression also appeared to be more dependent on H3K9 methylation than on NL tethering ^55^.

### Limitations of the study

*Only one locus investigated.* While our *loxP*-SB system greatly facilitates the generation of series of deletions and inversions in a locus of interest, it is not easily scalable to multiple loci. As a result, our study focused on one locus of about 10 Mb in the mouse genome. Further experiments will be needed to expand our results genome-wide. While we identified boosting regions in the two dissected LADs, the proportion of LADs that use this mechanism for their NL association is still to be determined.

*Only one cell type investigated.* Similarly, our analyses are so far restricted to F1 hybrid mESCs. Upon cell differentiation, ∼30% of the genome switches between LAD and iLAD classification ^9^ and lamina composition is also remodelled ^64,67^. This suggests that LAD-NL contacts are mediated to some extent by cell type specific mechanisms. A full comprehension of LAD formation will require their investigation in various cell types and even across species.

*Molecular foundation of boosting regions remains to be determined.* Our study was aimed to reveal overall principles of LAD - NL interactions. We did not fine-map possible NL - tethering sequence elements within the boosting regions. It remains to be determined whether such fine-mapping is feasible, or whether the boosting regions harbour a multitude of elements that are scahered throughout. More generally, the molecular mechanisms that are responsible for the affinity of boosting regions for the NL remain to be elucidated.

## METHODS

### Experimental procedures

#### Cell culture

F121/9 (CAST/EiJ x S129/Sv) female mouse embryonic stem cells (mESCs) F1 hybrid cell line ^68^ and derived clones were cultured in 2i+LIF condition, according to the 4D Nucleome guidelines (https://data.4dnucleome.org/biosources/4DNSRMG5APUM/). Briefly, cells were cultured in serum-free DMEM/F12 (Gibco) and Neurobasal (Gibco) medium (1:1), supplemented with N-2 (Gibco), B-27 (Gibco), 0.05% BSA (Gibco), 10 × 4 U leukemia inhibitory factor (LIF; Millipore), 1 μM MEK inhibitor PD0325901 (Mirdametinib, MedChemExpress), 3 μM GSK-3β inhibitor CHIR99021 (Laduviglusib, MedChemExpress), 1.5 × 10−4 M 1-thioglycerol (Sigma-Aldrich), and 1% penicillin-streptomycin (Gibco, 15070063) on 0.1% gelatin-coated plates. Cells were passaged every 2 days. Cells were seeded and incubated overnight before transfection for SB hopping, Cre recombination, CRISPR/Cas9 editing, as well as pA-DamID and RNA-seq experiments. Mycoplasma contamination was ruled out by regular testing (#LT07-318; Lonza).

### Insertion of the loxP-SB cassette

#### Transfection of the loxP-HyTK cassette

We cloned the vector pLD036 containing a *loxP* site and the double selection marker HyTK flanked by both FRT and F3 flipping sites and SB 3’ and 5’ inverted terminal repeats (ITRs). This cassette was amplified by PCR and knocked-in in the genome using CRISPR/Cas9 editing. We designed guide RNAs (gRNAs) targeting an intergenic sequence close to LAD1 5’ border using CHOPCHOP ^69,70^ and assessed their cutting efficiency by TIDE ^71^. The best gRNA was selected and assembled into an RNP together with the AltR CRISPR tracrRNA (1073189, IDT) and the Cas9 nuclease (1081058, IDT). We transfected 300,000 cells with the RNP and the PCR products using lipofectamine CRISPRMAX Cas9 Transfection Reagent (CMAX00001, Thermofisher Scientific) according to the manufacturer’s protocol. Three days after transfection, Hygromycin was added to the medium (200 µg/mL, 10687010, Invitrogen) to select for HyTK-expressing cells. We picked colonies six days later and screened clones for cassette integration by PCR and Sanger sequencing.

#### loxP swapping

We cloned the vector pLD037 containing a *loxP* site flanked by FRT and F3 flipping sites. We used Recombinase-Mediated Cassehe Exchange to flip the HyTK gene for a *loxP* site in the previously selected clone. We transfected 300,000 cells with 1.5 µg of flippase-encoding plasmid (Addgene #13787) and 0.5 µg of pLD037 using lipofectmamine 2000 (Invitrogen, 11668019). Two days after transfection Ganciclovir was added to the medium (2,5 µg/mL) to select for HyTK non-expressing cells. Colonies were picked seven days later and screened by PCR and Sanger sequencing.

##### SB hopping and SB hopped clone selection

For every hopping experiment, one million cells were transfected with 4.5 µg of pME07, a bicistronic vector encoding for the SB transposase SB100x and the human nerve growth factor receptor (LNGFR). We used lipofectamine 2000 according to the manufacturer’s protocol. To select for transfected cells expressing the surface marker LNGFR, ∼30h after transfection cells were submitted to MACS column selection using MS columns (130-042-201, Miltenyi Biotec) and LNGFR MicroBeads (130-091-330, Miltenyi Biotec). Non transfected cells were taken along as a negative control. Cells were grown for one week and single cells were sorted to grow clones. Crude extracts were prepared for each clone: cells from a full 96-plate well were lysed in 25 µL DirecPCR Lysis Reagent (302-C, Viagen Biotec) supplemented with 100 µg/mL proteinase K and incubated at 55°C, 2h30 and at 85°C, 45 min. In experiments in which SB hopped from the *loxP*-SB cassette, SB launch site was amplified by PCR using 2-3 µL of crude extracts. A shorter band was amplified for hopped clones compared to non-hopped ones. In experiments in which SB hopped from either LAD1 or LAD2, the junction between one arm of SB and the donor site was amplified by PCR. A band was amplified in non-hopped clones only. Over all experiments, SB hopped in 34% of the clones on average. The clone used to hop SB within LAD1 (**Figure 1D**) has two SB integrations. The extra SB copy is located 28 Mb away from LAD1 on chromosome 9. Data not shown were filtered out.

##### Mapping SB integration sites by Tagmentation

To map SB integration site after hopping, we adapted the Tagmentation protocol from ^72^ to amplify the junction between SB ITRs and genomic DNA. The overall procedure was similar to the one described in ^72^ except for the primers used in the PCR reactions. The linear enrichment PCR amplification was performed using primers MEP9 (5ʹ ITR) or LD027 (3ʹ ITR). PCR1 was done with primers MEP11 (5ʹ ITR) or MEP34 (3ʹ ITR) while we used primer Ib569 for PCR2 (both ITRs). SB integrations were validated by PCR in clones of interest.

##### Cre recombination

One million cells were transfected with 4 µg of vector encoding for the Cre recombinase fused to the fluorescent protein CFP, using lipofectamine 2000. The following day CFP positive cells were sorted by FACS and grown for a week. After that cells were sorted again, as single cells, to grow clones. To screen for recombined clones, crude extracts were prepared (see SB hopping section) and three PCR reactions were set-up for each clone. The procedure is illustrated in **Figure S11**. In brief, the anchor *loxP* site is flanked by 3’ and 5’ exogenous sequences after SB hopping while the hopping *loxP* site is flanked by SB 3’ and 5’ ITRs. After recombination, *loxP* sites are flanked by either the 5’ exogenous sequence and the 5’ SB ITR or by the 3’ exogenous sequence and the 3’ SB ITR. A clone is considered as recombined if its *loxP* sites are no longer flanked by both exogenous sequences or both SB ITRs but are flanked by one exogenous sequence (3’ or 5’) and one SB ITR (3’ or 5’ respectively). For deletion clones, we also made sure that the deleted piece could not be amplified from the genome. The hopped *loxP* site flanked by both SB ITRs are amplified in PCR1 with primers o-JOYC222 and o-JOYC186. *loxP* sites flanked by both exogenous sequences and the recombined junction between SB 3’ ITR and the 3’ exogenous sequence are amplified in PCR2, with primers o- JOYC187 and o-LD022. Finally, the recombined junction between the 5’ exogenous sequence and the SB 5’ ITR are amplified in PCR3 with primers o-LD087 and o-LD100. All three PCR reactions contain 10 µL of MyTaq HS Red mix (BIO-25048, Bioline), 2-3 µL of the crude extract and 1 µL of each primer (10 µM) for a final volume of 20 µL. Cre recombination efficiency was measured as the number of recombined clones over the total number of screened clones.

##### R4-5 CRISPR deletion

We deleted R4-5 in a mESCs cell line expressing the ddCas9 construct from the ROSA28 locus. We used CHOPCHOP ^69,70^ to design CAST allele-specific guide RNAs. Pools of five and four guide RNAs were designed to target R4-5 5’ and 3’ end respectively. gRNAs were cloned into a mCherry expressing plasmid and pooled. One million cells were transfected with 4 µg of the pool of gRNAs using lipofectamine 2000. Shield-1 (500 nM, AOB1848, Aeobius) was added to the medium to stabilize the ddCas9 protein. The day after, mCherry positive cells were sorted by FACS and kept in culture for a week in Shield-containing medium. Single cells were sorted, clones were expanded and screened by PCR and Sanger sequencing.

##### R4-5 and R2-3 isolation in a strong iLAD and close to LAD4

We used CHOPCHOP ^69,70^ to design CAST allele-specific gRNAs. gRNAs targeting iLAD regions were cloned in a CFP expressing plasmid while those targeting LAD1 were cloned in an mCherry expressing plasmid. One million cells were transfected with the corresponding pools of guide RNAs and with the pX458 plasmid encoding for Cas9 and GFP proteins (48138, Addgene), using lipofectamine 2000. The day after, triple positive cells (GFP+, mCherry+, CFP+) were sorted by FACS and kept in culture for a week. After that, triple negative cells (GFP-, mCherry-, CFP-) were sorted as single cells, clones were expanded and screened by PCR and Sanger sequencing.

##### pA-DamID (LaminB1 and H3K9me3)

The library preparation was performed as follows: genomic DNA was isolated (Bioline, bio-52067) and ∼ 500 ng were digested with DpnI (10 U, NEB, R0176L) in CutSmart Buffer 1X (8h 37°C, 20 min 80°C) in a total volume of 10 µL. A-tailing was performed by addition of 5 µL of the A-tailing mix (0.5 µL of Cutsmart buffer 10X, 0.25 µL Klenow 50 U/µL (NEB, M0212M), 0.05 µL dATP 100 mM, 4.2 µL H2O) and incubation 30 min at 37°C followed by 20 min at 75°C. Adapters were ligated by adding 15 µL of the ligation mix (3 µL T4 Ligase Buffer 10X, 0.5 µL T4 ligase (5 U/µL, Roche, 10799009001), 0.25 µl of x-Gene Stubby Adapter 50 mM (IDT), 11.25 µl H2O) and incubating samples 16h at 16°C and 10 min at 65°C. Finally, the Methyl Indexed PCR was performed by mixing 4 µL of ligated DNA with x-Gen Dual combinatorial Indexes (IDT) (125 nM final concentration) and MyTaq RedMix (Bioline, BIO-25048) in a final volume of 40 µL. The following PCR program was used: 1 X (1 min 94°C), 14 X (30 sec 94°C, 30 sec 58°C, 30 sec 72°C), 1X (2 min 72°C). The resulting amplified material was processed for high-throughput sequencing and sequenced for single-end 100-bp reads on a NovaSeq 6000 platform. Approximately 30 million reads were sequenced for every condition.

For LaminB1 pA-DamID experiments, we processed two independent recombined or CRISPR/Cas9 edited clones except for the isolation of R2-3 in an iLAD and close to LAD4 where only one edited clone could be selected. For H3K9me3 pA-DamID experiments, we processed one recombined clone for recombination d3, d6, i10 and i12 and two recombined clones for recombination i7.

##### RNA-seq

One million cells were harvested, washed once in cold PBS, resuspended in 600 µL of RLT buffer (RNeasy Mini kit, 74104, Qiagen) and stored at -80°C until subsequent RNA isolation. RNA was isolated using the RNeasy Mini kit (74104, Qiagen). Libraries were prepared using the TruSeq® Stranded mRNA Library Prep (20020594, Illumina) and TruSeq RNA Single Indexes Set A (Illumina) kits. Libraries were sequenced for single-end 75-bp reads on a NextSeq 550 platform. Approximately 25 million reads were sequenced for every condition.

For recombination d6, i10 and i12 one recombined clone was processed while two recombined clones were processed for recombination d3 and i7.

### Computational Analysis

#### Tagmentation mapping

We adapted the pipeline published in ^74^ and created a build-in R package to improve user usage. The pipeline was modified to map integration sites of the SB transposon and determine if the integration occurred on the CAST or on the 129S1 allele.

#### pA-DamID

To process pA-DamID data, we adapted a previously published pipeline ^73^, to align reads to the mouse reference genome (Release M23 – mm10) ^75^ and map them to the CAST and 129S1 alleles. We used the WASP toolkit ^76^ for the latter step, providing SNPs data from CAST and 129S1 alleles, available on the Mouse Genomes Project (https://www.mousegenomes.org/publications/) ^77,78^.

LAD coordinates were determined using hidden Markov modeling on the average NL interaction profile between biological replicates, in wild-type (WT) F1 hybrid mESCs (https://github.com/gui11aume/HMMt).

Differential pA-DamID tracks of NL interactions were smoothed using the “runmean” function from the caTools package, using a moving window of width 7 (k = 7).

#### Domainograms

To visualize changes in NL interactions, we adapted a previously described pipeline ^41^, to analyse genomic deletions and inversions. For genomic deletions: pA-DamID score computed for each GATC fragment of the non-recombined genomes were smoothed. For recombined genomes, GATC fragments overlapping the deleted region were removed before smoothing. pA-DamID plots and domainograms are plotted on the reference genome. For genomic inversions: coordinates of the GATC fragments overlapping the inverted region were inverted and data was then smoothed. Because pA-DamID plots and domainograms are plotted on the inverted genome, the same GATC fragments were inverted after smoothing for the non-recombined genomes. The pipeline is available at: https://github.com/vansteensellab/domainograms/blob/dauban_etal/plot_domainograms.R.

For combined inversions and CRISPR/Cas9 deletions, the coordinates of GATC fragments overlapping the inverted region were inverted in-silico, then the GATC fragment overlapping the deleted region were removed and data were smoothed. The pipeline is available at: https://github.com/vansteensellab/domainograms/blob/dauban_etal/plot_domainograms_combined_recombination.R.

All pA-DamID scores were converted to z-scores to minimize variability across experiments and obtain comparable dynamic ranges. Across all experiments, one z-score unit corresponds on average to 0,87 log2-unit.

Gene expression was computed from publicly available RNA-seq experiments performed in WT F1 hybrid mESCs (4DNFIPFKK5LM). Data are not allele-specific.

The CAST (resp. 129S1) allele in the recombined clone is always plotted against the CAST allele (resp. 129S1) in the non-recombined one.

#### Transcription factor motif scanning

The number of cKrox, YY1 and Trl motifs in R4-5, the remainder of LAD1, LAD2.1 and LAD2.2 was computed using MotifScan (https://motifscan.readthedocs.io/en/latest/index.html). The weight matrices of cKrox and YY1 motifs were downloaded from the latest version of the Hocomoco database (ZBT7B.H12CORE.0.SM.B and H12CORE.0.PSM.A, respectively). The weight matrix of Trl motif was downloaded from the JASPAR database (MA0205.1) ^79^. The number of motifs detected in each region was then normalised to the regions’ respective size.

#### Allele-specific RNA-seq data processing

We obtained the 129S1 (GCA_001624185.1) and CAST/Eij (GCA_001624445.1) genome fasta files from the UCSC genome browser website (https://hgdownload.soe.ucsc.edu/hubs/mouseStrains/hubIndex.html). Reads were mapped to the two genomes separately (STAR aligner, ^80^) and only those with MAPQ > 10 were kept (SAMtools, ^81^). Reads coming from each allele were then split using a custom R script. Briefly, the alignment score (AS tag in BAM file) of a given read to the 129S1 genome is compared to its alignment score to the CAST genome. If it is higher for 129S1 genome (resp. CAST), the read is assigned to 129S1 genome (resp. CAST). If alignment scores are equal and greater than zero, the read is tagged as ambiguous. If both alignment scores are zero, the read is tagged as unmapped. Finally, split allele-specific and ambiguous/unmapped bam files are produced as output. To generate allele-specific gene counts, gene annotations from GENCODE (Release M23 – mm10) ^75^ were first lifted over to 129S1 and CAST genomes using Liftoff (https://github.com/agshumate/Liftoff) ^82^. Then, read coordinates from the allele-specific filtered bam files were overlapped with the corresponding lifted-over gene coordinates, using the “summarizeOverlaps” function from the GenomicAlignments Bioconductor package ^83^. These allele-specific read counts were used for subsequent analyses.

To assess gene expression changes, we first normalized read counts in the parental cell line over the total amount of reads in the non-recombined one. For each gene of interest, we then compared the number of reads coming from the CAST allele (resp. 129S1) in the recombined clone to the number of reads coming from the CAST allele (resp. 129S1) in the non-recombined clone, over the total amount of reads detected in each allele (resp. 129S1). We performed Fisher tests on these gene-specific matrices and p-values were corrected for multi-testing by multiplying them by the number of tests performed. Genes were considered as significantly down or up-regulated if the Fisher test p-value was lower than 0.05 and the fold change above two.

In recombination d3 and d6, no CAST allele-specific reads were detected for most of the genes within the deleted region. A small number of CAST-specific reads were still detected on the Tdgf1 gene (with a 10-fold decrease in gene expression). These reads were discarded after visual inspection as they resulted from sequencing and mapping issues.

**Table S1.**
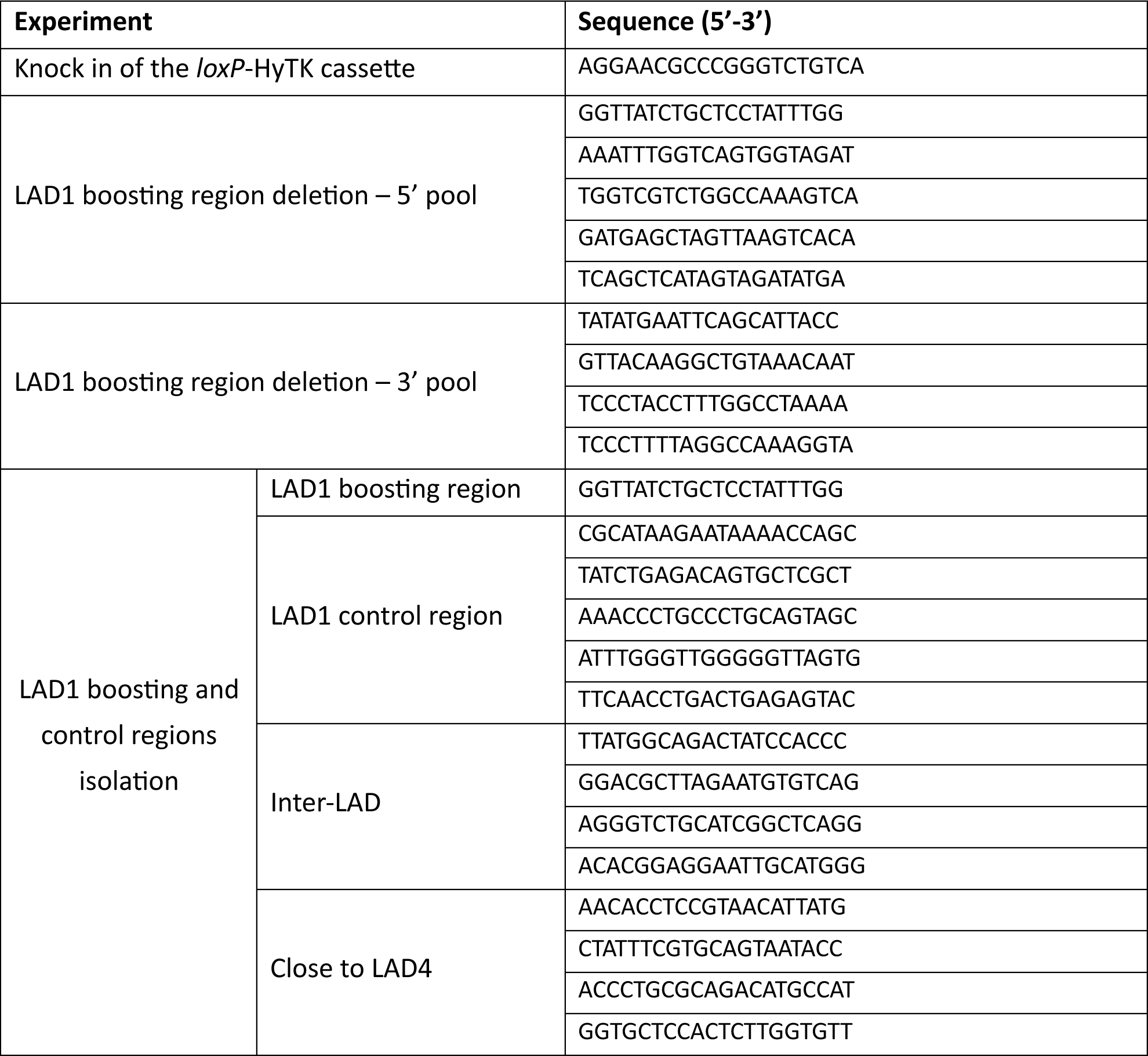
Lists of guide RNAs used in this study.

**Table S2.**
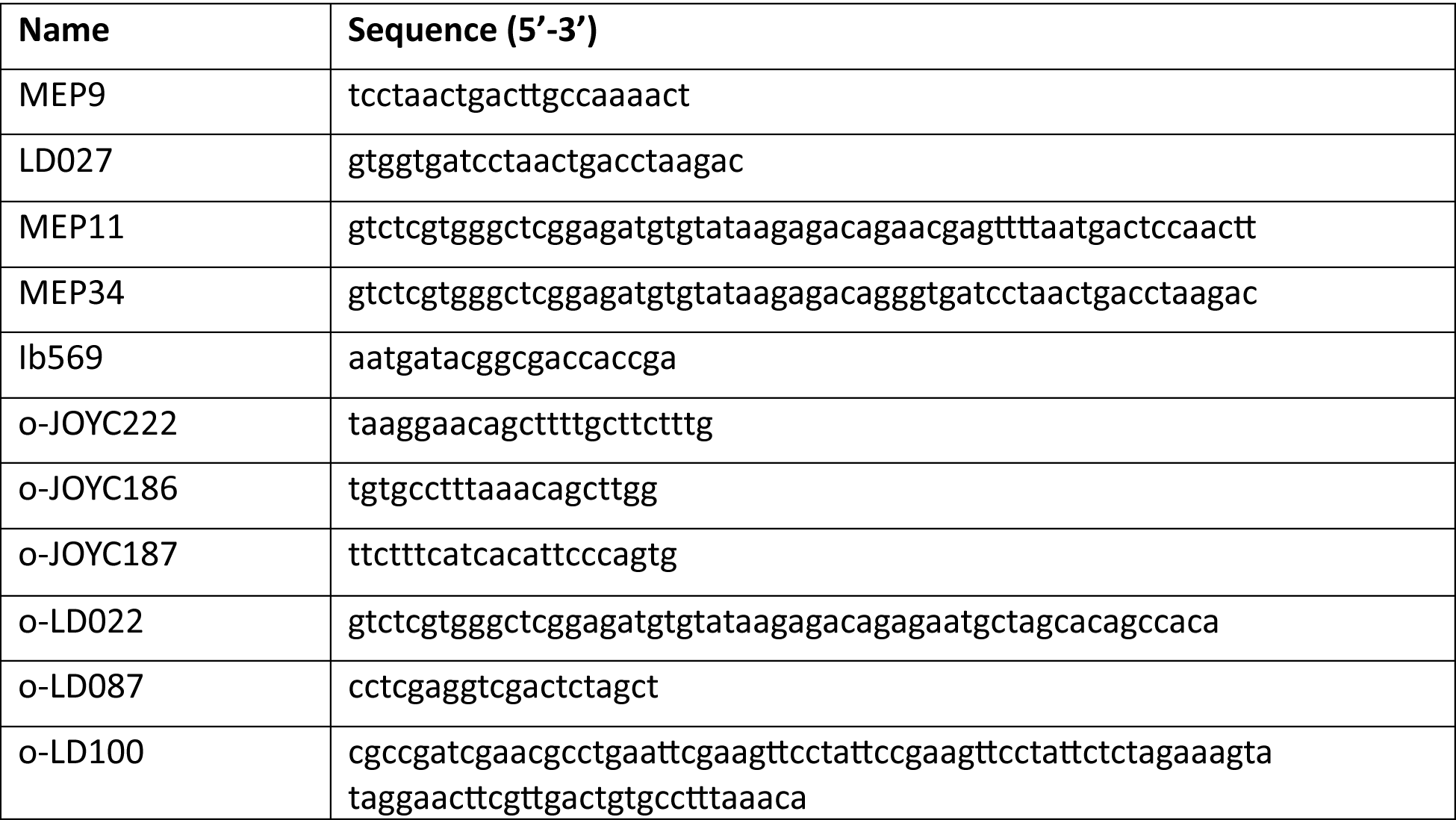
Lists of primers used in this study.

## DATA AVAILABILITY

Laboratory notebooks and supplementary data are available on Zenodo (10.5281/zenodo.10135191). Raw sequencing data and processed data are available on GEO under the accession number GSE250307. Code and supplementary files used as input for the scripts are available on Zenodo (10.5281/zenodo.10222688).

## ACKNOWLEDGEMENTS

We thank the NKI Genomics, Flow Cytometry, Protein Production, and Research High Performance Computing core facilities for excellent support, and members from our laboratories for inspiring and helpful discussions.

## FUNDING

This project has received funding from the European Union’s Horizon 2020 research and innovation programme under the Marie Sklodowska-Curie grant agreement No 101023923. It was also funded by the European Research Council: GoCADiSC, 694466 (B.v.S.); RE_LOCATE, 101054449 (B.v.S.); FuncDis3D, 865459 (M.M) and EMBO Long-term Fellowship (J.O.Y.C). E.d.W is supported by a Vici grant (VI.C.222.049) from the Dutch Research Council. Views and opinions expressed are however those of the author(s) only and do not necessarily reflect those of the European Union or the European Research Council. Neither the European Union nor the granting authority can be held responsible for them. Research at the Netherlands Cancer Institute is supported by an institutional grant of the Dutch Cancer Society and of the Dutch Ministry of Health, Welfare and Sport. The Oncode Institute is partially funded by the Dutch Cancer Society.

## AUTHOR CONTRIBUTIONS

Conceptualization: L.D., J.O.Y.C., B.v.S.

Wet-lab experiments: L.D., M.E., M.d.H., J.O.Y.C., M.M.A.

Data analysis and visualization: L.D., V.H.F.S.,

Method development: L.D., M.E., J.O.Y.C.

Code development: T.v.S., C.L., K.R., V.H.F.S., M.M.

Writing: L.D., B.v.S.

Funding Acquisition: B.v.S., L.D.

Supervision: B.v.S.

## DECLARATION OF INTERESTS

The authors declare no conflict of interest.

---

**Figure S1:**
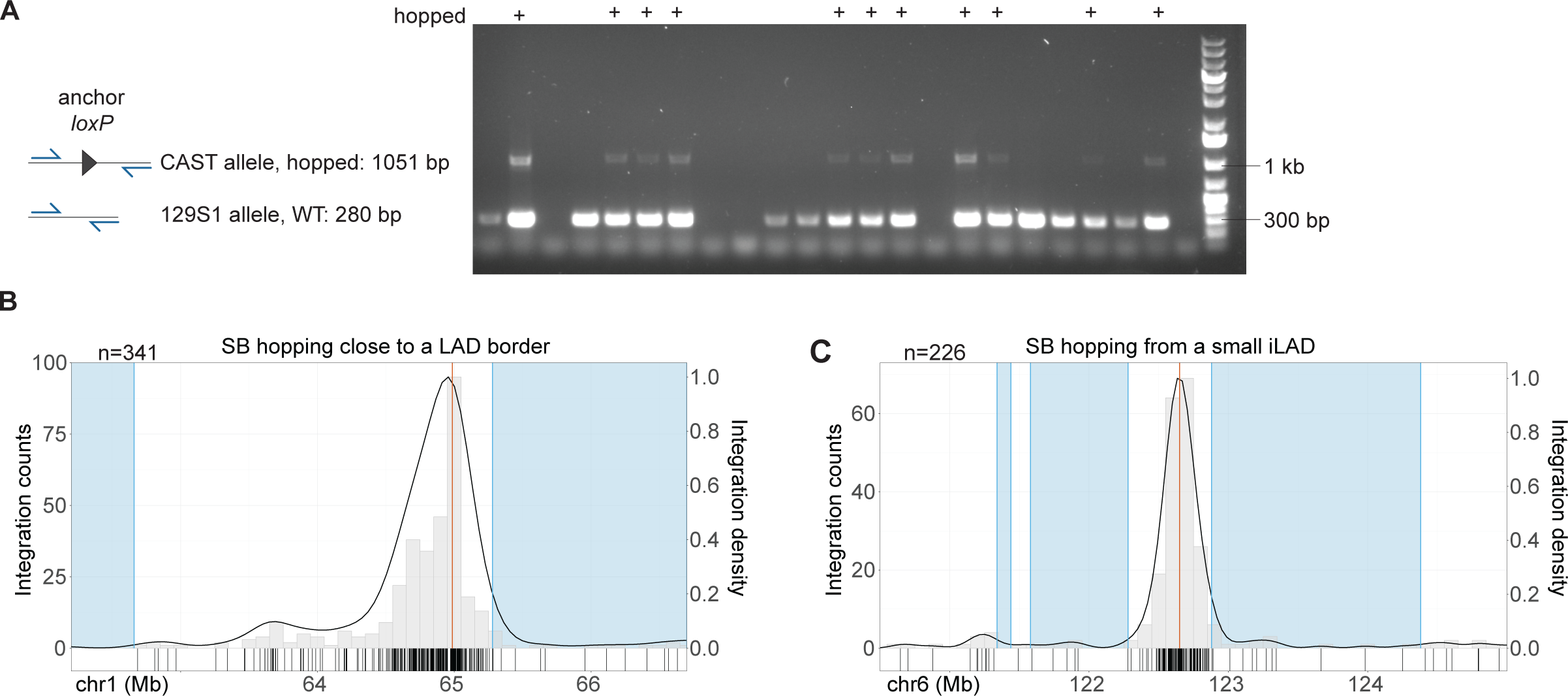
Screening for clones with hopped SB; when hopped from an iLAD launch site, SB remains mostly within the iLAD. **(A)** PCR screening of clonal cell lines obtained after mobilization of *loxP*-SB from the launch site located on the CAST allele in the iLAD between LAD1 and LAD2 (see Figure 1A). A 129S1 allele-specific band is expected to be amplified in all clones. In SB hopped clones an additional product is amplified from the CAST allele, including the anchor *loxP*. In the absence of hopping, the CAST allele is expected to yield a PCR product of 1872 bp; however, this product is inefficiently amplified due to its larger size. **(B, C)** Distribution of SB integrations after hopping from launch sites close to a LAD border (B) or in a small iLAD (C). The histogram of the number of SB integrations per 100kb bin (left y-axis) is overlaid with the density curve (right y-axis). Each tick on the x-axis represents a unique SB integration event. SB integrations were mapped both from cell pools and clonal cell lines. n corresponds to the number of unique SB integrations mapped in the plotted windows. Red line: launch site, blue rectangles: LADs.

**Figure S2:**
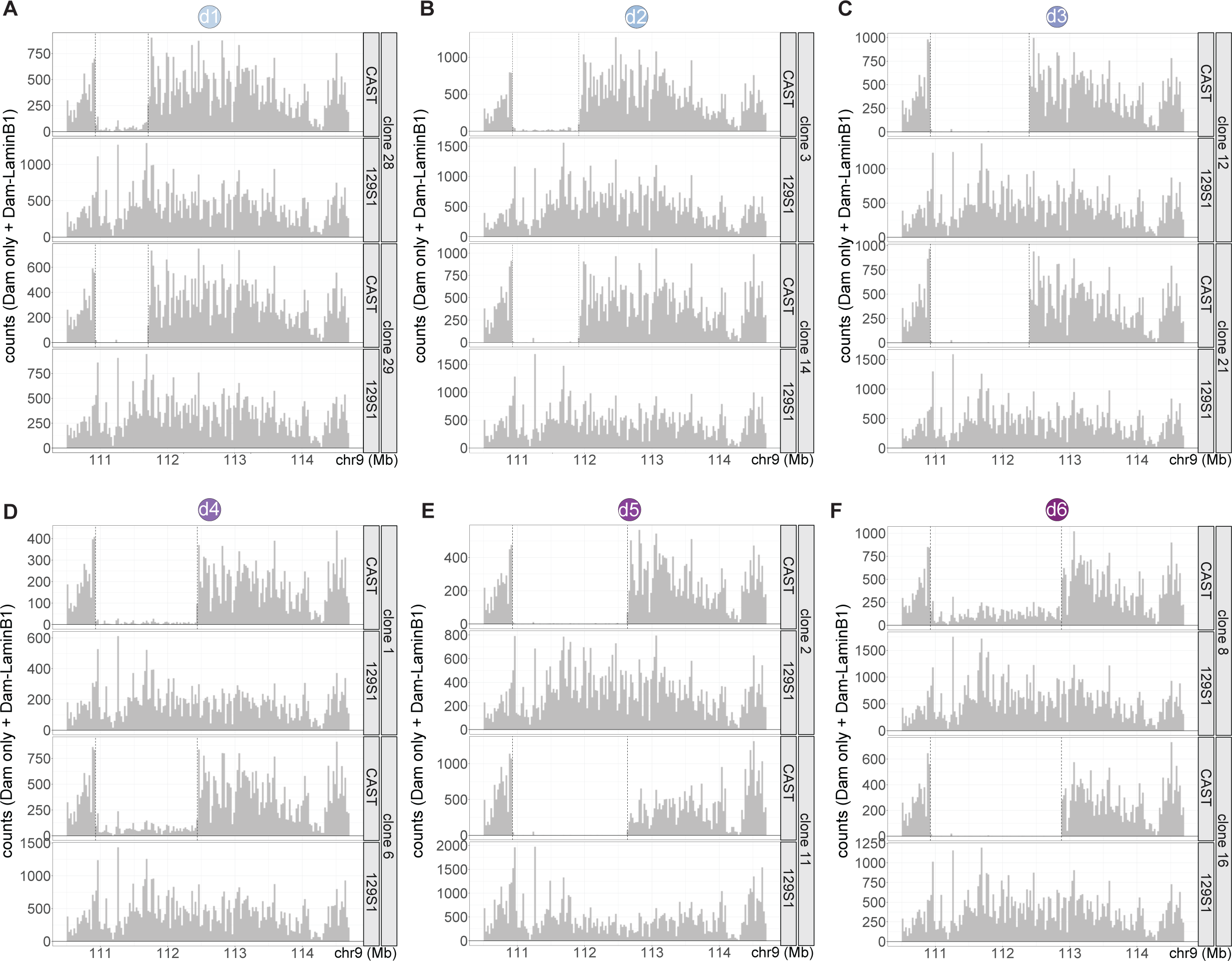
Validation of CAST-allele deletions. Total CAST and 129S1 allele-specific sequencing read counts per 25 kb from pA-DamID experiments, at the recombination sites and flanking regions, for all recombined clones used in the deletion series d1 through d6. Read counts are the sum of Dam-LaminB1 and Dam-only reads; note that this is different from the pA-DamID tracks in Figure 2, which show the (z-score transformed) log2 ratio of Dam-LaminB1 and Dam-only. Vertical dashed lines indicate the expected recombination sites on the CAST allele. Note that a few reads are detected within the deleted region in clones from recombination d1 (A, clone 28), d4 (D, clone 6) and d6 (F, clone 8). These cell lines may consist of recombined cells with a low level of contamination from non-recombined cells.

**Figure S3:**
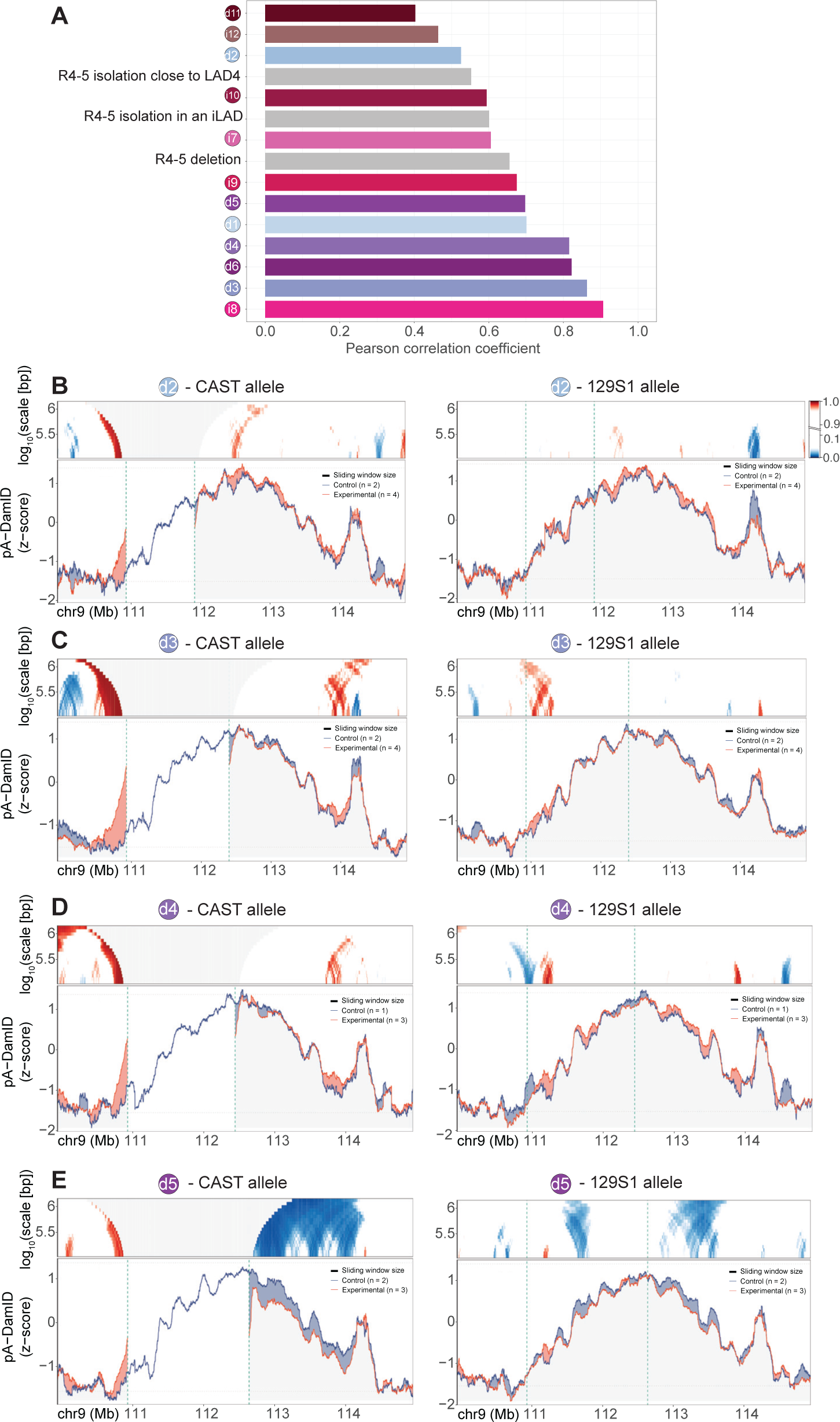
Reproducibility between independent clones for all tested rearrangements; additional data of deletion series. **(A)** Correlations between the two independent recombined or CRISPR/Cas9-edited clones for each tested rearrangement, calculated as the Pearson correlation coefficient of the changes in pA-DamID score (Experimental minus Control), for windows of 123 kb. **(B-E)** Same plots as in Figure 2B-C for recombination d2 (B), d3 (C), d4 (D) and d5 (E). Gene annotation tracks are not displayed. Domainogram colour-key in (B) applies to all panels.

**Figure S4:**
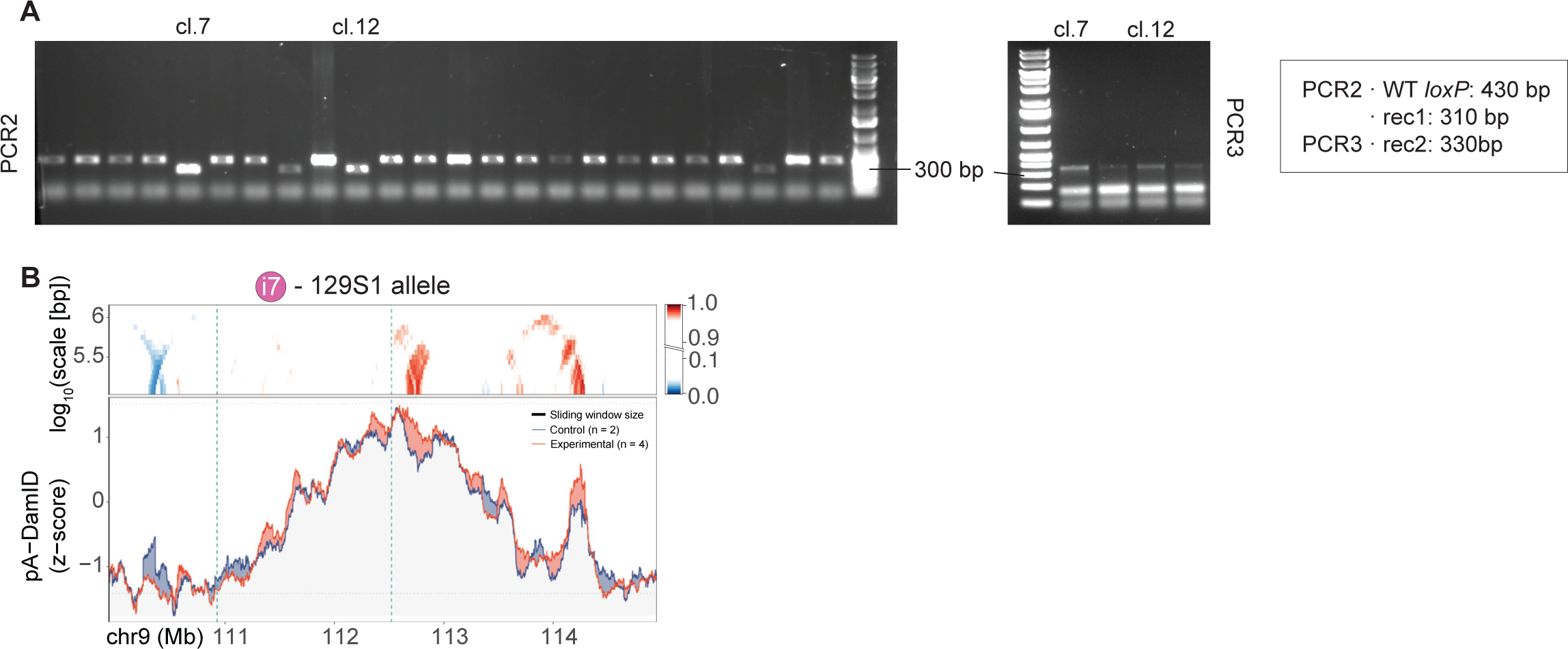
Validation of recombination i7 recombined clones. **(A)** Screening of clones from recombination i7 by PCR2 (left panel) and further analysis of candidate clones by PCR3 (right panel). For details of primers and expected PCR products, see Figure S11. Clones were selected if rec1 and rec2 products only were amplified by PCR2 and PCR3 respectively. PCR1 was not informative because these clones carry an extra copy of SB. This extra copy is located 28 Mb away on chromosome 9 and does not affect the current results. We chose clones 7 and 12 for pA-DamID experiments. **(B)** Same plot as in Figure 3B, for the 129S1 allele. pA-DamID tracks and domainograms are plotted on the WT chromosomal coordinates.

**Figure S5:**
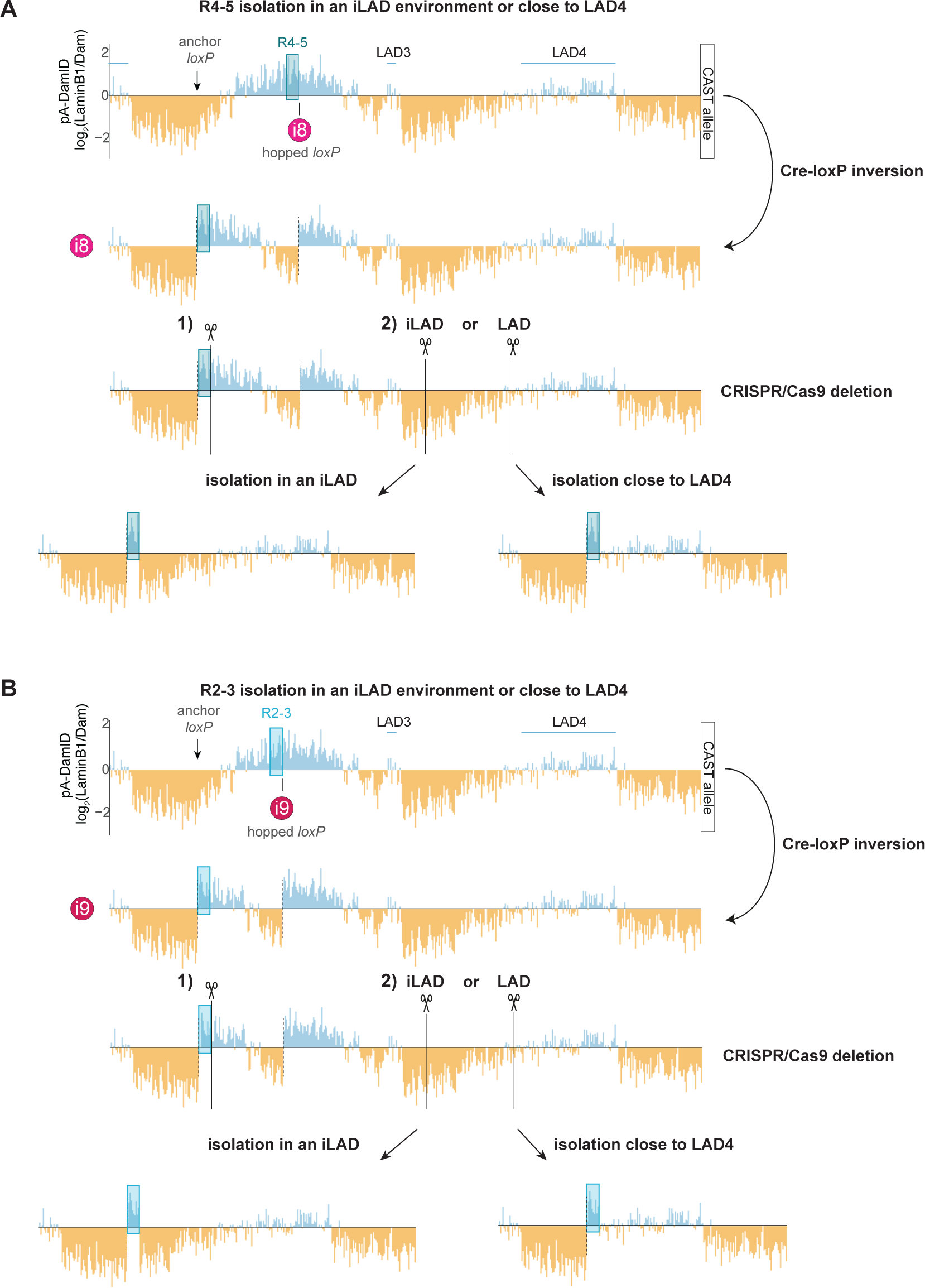
Strategy to relocate R4-5 and R2-3 to an iLAD environment or close to LAD4. **(A)** Simulated NL interaction tracks to illustrate two-step relocation of R4-5. First, upon recombination i8, R4-5 is flanked on one side by iLAD sequences. Second, to totally isolate R4-5 in an iLAD environment, the remaining LAD1 sequences and part of the flanking iLAD were deleted by two Cas9-mediated cuts in the CAST allele of recombined clones: at the R4-5-LAD1 junction and in the middle of the following iLAD. To isolate R4-5 close to LAD4, the second cut was induced closer to LAD4. **(B)** Simulated NL interaction tracks to illustrate two-step relocation of R2-3. Upon recombination i9, R2-3 is flanked on one side by iLAD sequences. Next, to isolate R2-3 in an iLAD or close to LAD4, we used the same strategy as in (A).

**Figure S6:**
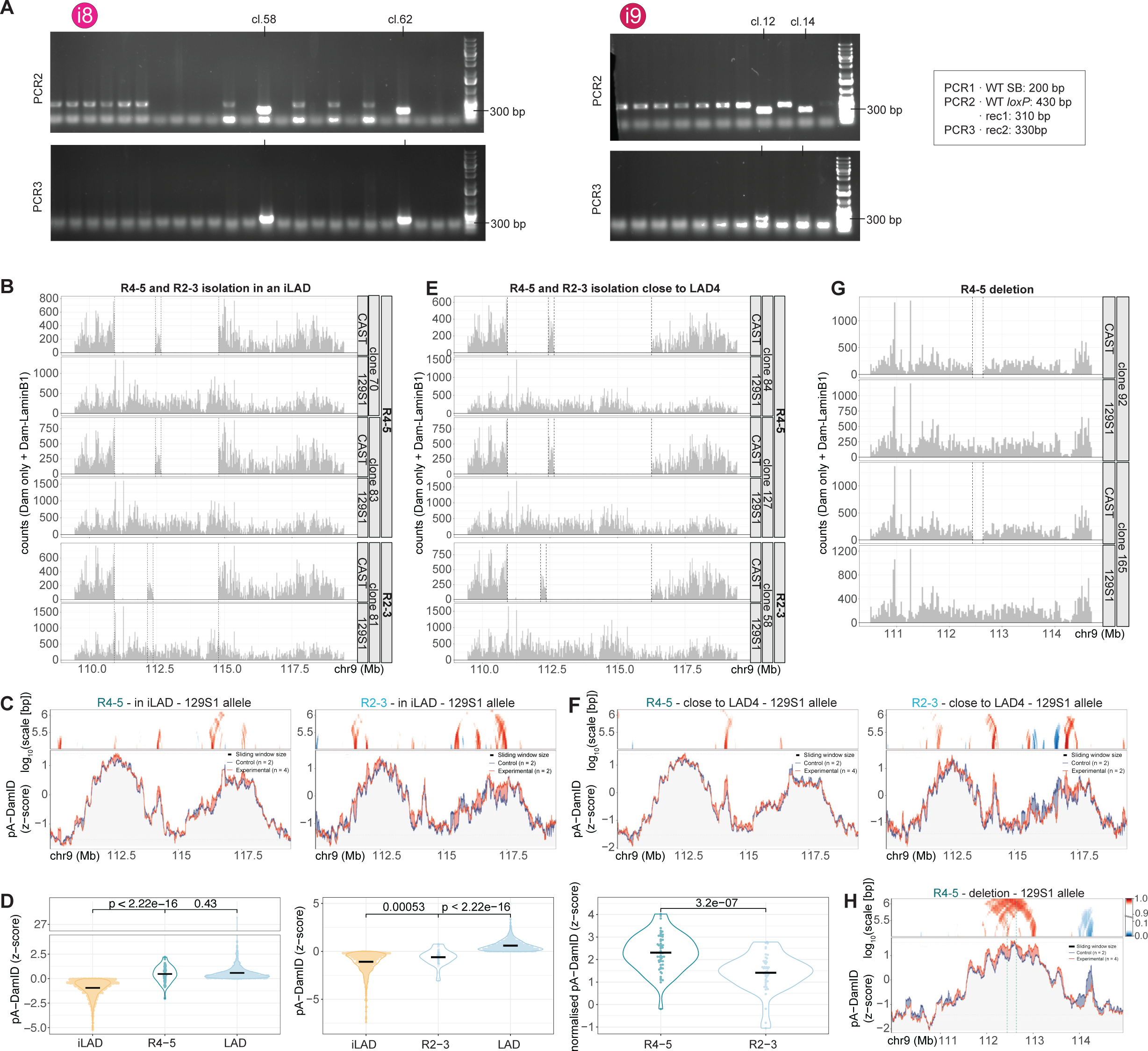
Screening, validation and additional analysis of R4-5 and R2-3 relocation clones. **(A)** Screening of clones from recombination i8 and i9 by PCR. Clones were selected if rec1 and rec2 products only were amplified by PCR2 and PCR3 respectively. For PCR design, see Figure S11. PCR1 was not informative because these clones carry an extra copy of SB. **(B, E)** Bar plots showing the total CAST and 129S1 allele-specific sequencing read counts per 25 kb from pA-DamID experiments, at the Cas9-mediated deletion sites and flanking regions, in clones in which R4-5 (top) or R2-3 (bottom) were isolated in an iLAD environment (B) or close to LAD4 (E). Read counts are the *sum* of Dam-LaminB1 and Dam-only reads; note that this is different from the pA-DamID tracks in Figure 2, which show the (z-score transformed) *log_2_ ratio* of Dam-LaminB1 and Dam-only. Vertical dashed lines indicate the expected deletions on the CAST allele. **(C, F)** Same plots as in Figure 3D (C) and Figure 3F (F), but for the 129S1 allele. pA-DamID tracks and domainograms are plotted using WT chromosomal coordinates. Domainogram colour-key is displayed in (H). **(D)** Left panel: R4-5 contacts the NL as frequently as its neighbouring LAD. pA-DamID score (z-score) for NL interactions of iLAD, R4-5 or LAD GATC fragments from the window plotted in Figure 3D, left. Middle panel: R2-3 is not as strongly associated with the NL as its neighbouring LAD. Same but for iLAD, R2-3 or LAD GATC fragments from the window plotted in Figure 3D, right. Right panel: R4-5 has a higher NL contact frequency than R2-3. For these analyses, R4-5 and R2-3 GATC fragments more than 2.5 kb apart were normalised over the 5th percentile of genome-wide pA-DamID values and their normalised pA-DAmID score distributions plotted. Because the resolution of DamID is estimated to be about 1 kb, we conducted this analysis on a subset of GATC fragments that are at least 2.5 kb apart, thus ensuring independent measurements. Black lines show median values. Statistical significance was assessed using Wilcoxon tests. **(G)** CAST and 129S1 allele-specific pA-DamID read counts (sum of Dam-only plus Dam-LaminB1 reads) in clones in which R4-5 was deleted. Vertical dashed lines indicate the **expected** deletion on the CAST allele. **(H)** Same plot as in Figure 3G, for the 129S1 allele. pA-DamID tracks and domainograms are plotted using WT chromosomal coordinates.

**Figure S7:**
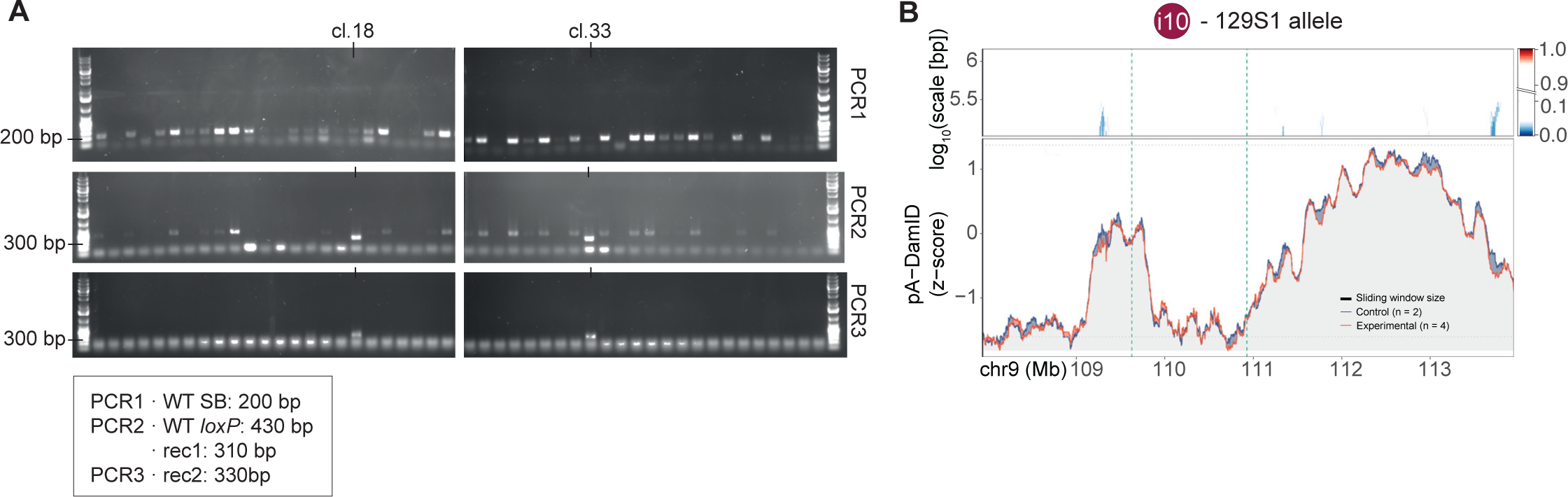
Validation of recombination i10 clones. **(A)** Screening of clones from recombination i10 by PCR. Clones were selected if PCR1 showed no amplification of the WT SB product and rec1 and rec2 products only were amplified from PCR2 and PCR3 respectively. For more details, see Figure S11. **(B)** Same plot as in Figure 4B, for the 129S1 allele. pA-DamID tracks and domainogram are plotted using WT chromosomal coordinates.

**Figure S8:**
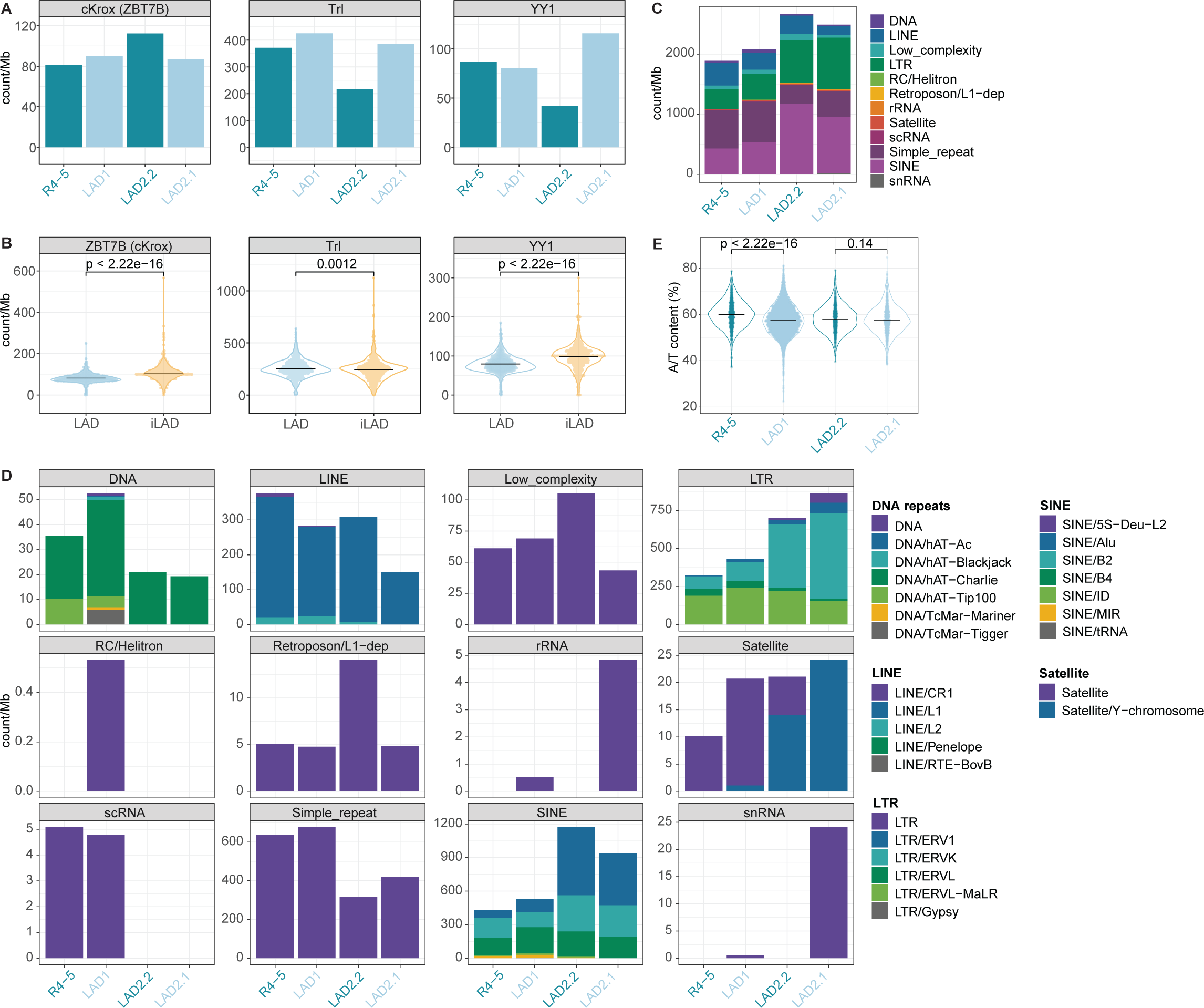
Sequence analysis of boosting regions. **(A)** Average density (motif count per Mb) of cKrox, Trl, and YY1 motifs in R2-3, R4-5, and LAD2.2. **(B)** Distributions of the number of cKrox, Trl, and YY1 motifs per Mb in LADs and iLADs. Black lines show median values. P-values according to Wilcoxon tests. **(C)** Number of repeats (according to RepeatMasker; https://www.repeatmasker.org) per Mb in R4-5, LAD1 lacking R4-5 sequences, LAD2.2, and LAD2.1. **(D)** Same as in (C) but for subclasses of repeat elements. **(E)** Percentage of A and T nucleotides computed in non-overlapping windows of 250 bp, in R4-5, LAD1 without R4-5, LAD2.2 and LAD2.1. Black lines show median values. P-values according to Wilcoxon tests.

**Figure S9:**
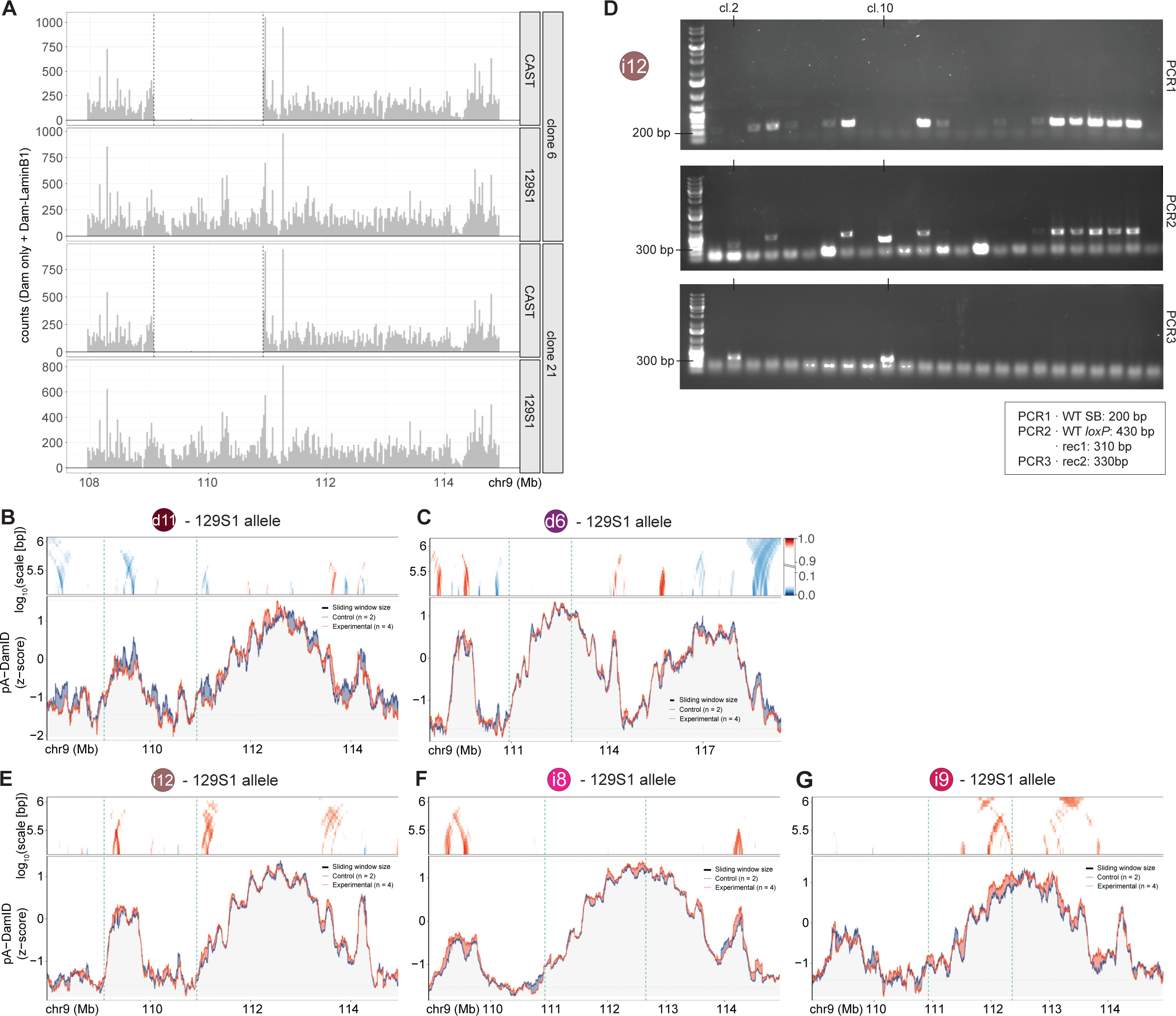
Validation and additional analyses of d11 and i12 recombined clones. **(A)** Total CAST and 129S1 allele-specific sequencing read counts per 25 kb from pA-DamID experiments, at the recombination sites and flanking regions, for clones used for deletion d11. Vertical dashed lines indicate the expected deletion on the CAST allele. **(B)** Same plot as in Figure 5B, for the 129S1 allele. pA-DamID tracks and domainograms are plotted using WT chromosomal coordinates. Domainogram colour-key is displayed in (C). **(C)** Same plot as in Figure 2C, right, for a wider window. **(D)** Screening of clones from recombination i12 by PCR. Clones were selected if PCR1 showed no amplification of the WT SB product and rec1 and rec2 products only were amplified from PCR2 and PCR3 respectively. For more details, see Figure S11. **(E-G)** Same plots as in Figure 5D (E), Figure 5E (F) and Figure 5F (G), for the 129S1 allele. pA-DamID tracks and domainograms are plotted on the WT chromosomal coordinates. Domainogram colour-key in (C) also applies to (E-G).

**Figure S10:**
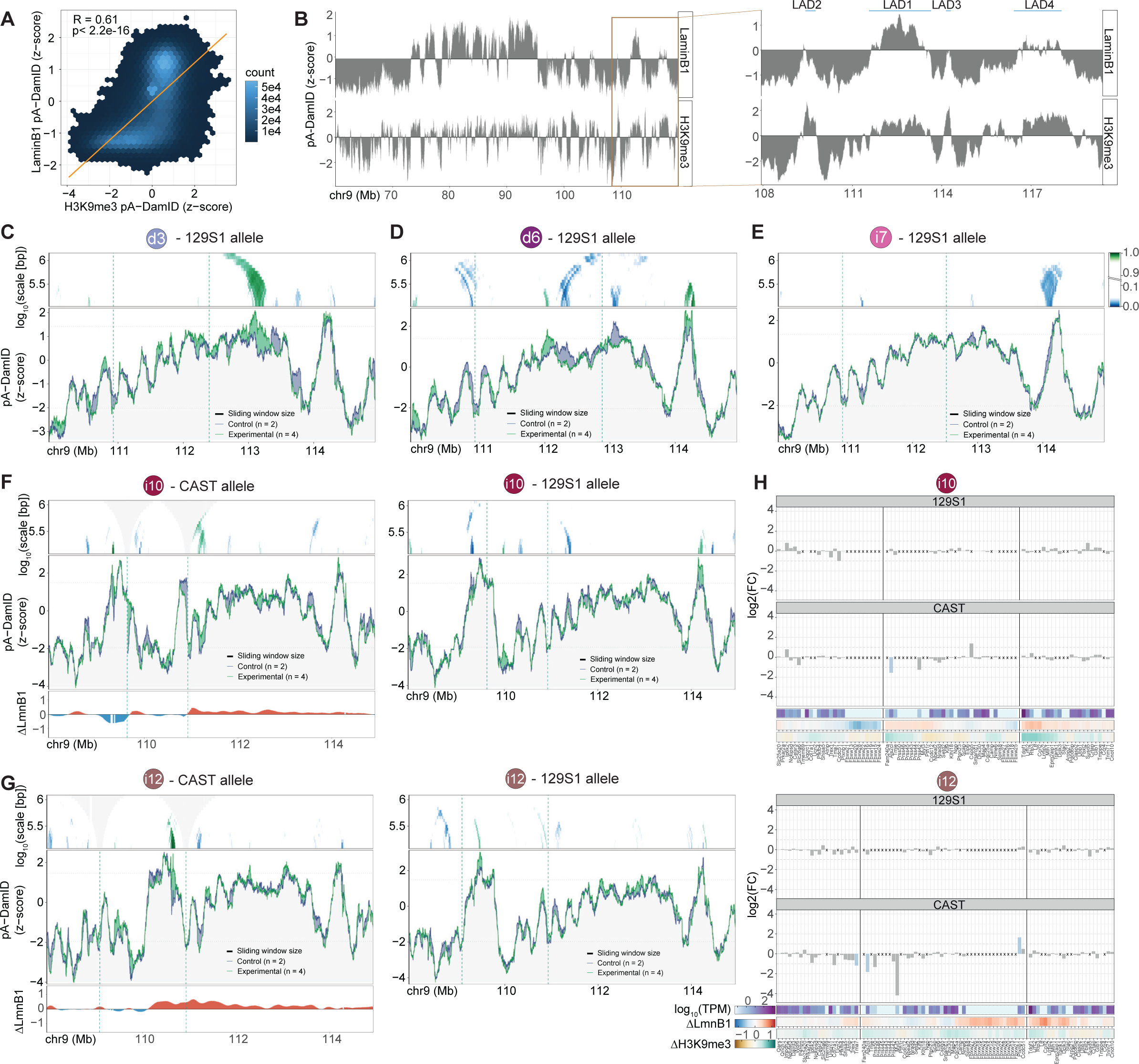
Spreading of NL contacts is not mirrored by changes in H3K9me3 and gene expression in recombination i10 and i12. **(A)** Genome-wide correlation between NL interactions (z-score) and H3K9me3 levels (z-score). Data were smoothed. Counts correspond to the number of GATC fragments per hexagonal bin. Orange line represents the linear fit of the data. R, Pearson’s correlation coefficient. **(B)** Smoothed pA-DamID tracks (z-score) of NL interactions (top) and H3K9me3 (bottom), of the 129S1 chromosome 9 in the parental clone of recombination i7. Left panel: overview of ∼60 Mb; brown box marks the region shown in close-up in the panel on the right, with positions of LAD1-LAD4 indicated. **(C-E)** Same plots as in Figure 6A (C), Figure 6B (D) and Figure 6C (E), for the 129S1 allele. pA-DamID tracks and domainograms are plotted using the WT chromosomal coordinates. **(F, G)** Left panel: Same plots as in Figure 6C for recombination i10 (F) and i12 (G). Right panel: same plot for the 129S1 allele. **(H)** Same plots as in Figure 6F for recombination i10 (top) and i12 (bottom).

**Figure S11:**
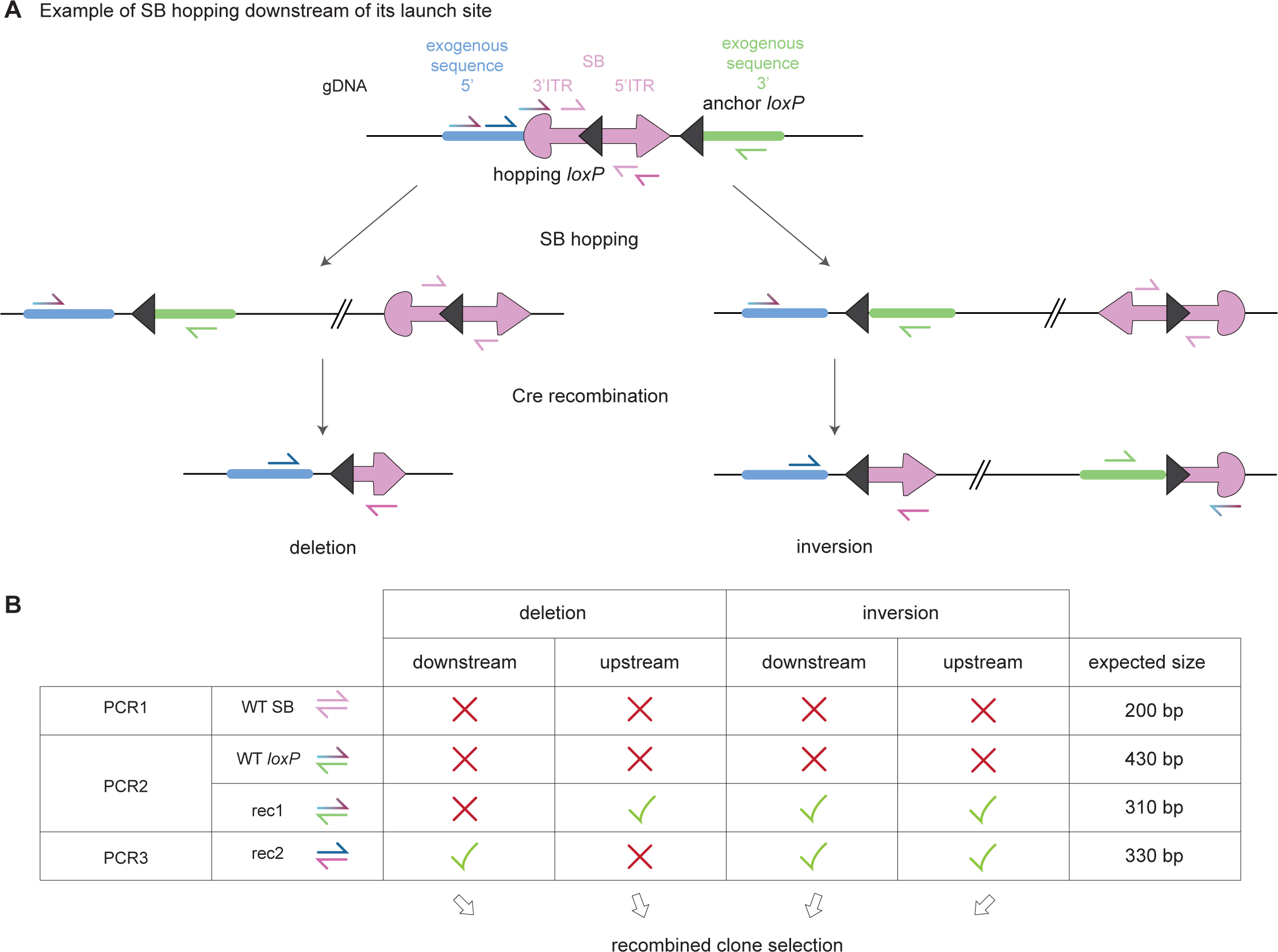
Schematic of rearrangements upon Cre recombination; clone selection by PCR. **(A)** Schematics of chromosomal re-arrangements induced by SB integration downstream of its launch site and subsequent Cre recombination. PCR primers used to characterise recombined clones are also depicted. **(B)** Table of primer pairs used to screen recombined clones and their expected output. PCR1 amplifies the WT, non-recombined SB (“WT SB” product). PCR2 amplifies (i) the anchor loxP prior to recombination (“WT loxP”) and (ii) the recombined junction between SB 3’ ITR and the 3’ region flanking the anchor loxP site (“rec1”). PCR3 amplifies the other recombined junction, between SB 5’ ITR and the 5’ region flanking the anchor loxP site (“rec2”). Recombined clones were selected if none of the WT products (“WT SB” or “WT loxP”) was amplified and either one (deletions) or both (inversions) recombined junctions were amplified.

## Notes

https://zenodo.org/records/10135191

https://zenodo.org/records/10222688

https://www.ncbi.nlm.nih.gov/geo/query/acc.cgi?acc=GSE250085

### Competing Interest Statement

The authors have declared no competing interest.

## References

1. Jerkovic, I., and Cavalli, G. (2021). Understanding 3D genome organization by multidisciplinary methods. Nat Rev Mol Cell Biol 22, 511–528. 10.1038/s41580-021-00362-w.

2. Rullens, P.M.J., and Kind, J. (2021). Attach and stretch: Emerging roles for genome-lamina contacts in shaping the 3D genome. Curr Opin Cell Biol 70, 51–57. 10.1016/j.ceb.2020.11.006.

3. van Steensel, B., and Belmont, A.S. (2017). Lamina-Associated Domains: Links with Chromosome Architecture, Heterochromatin, and Gene Repression. Cell 169, 780–791. 10.1016/j.cell.2017.04.022.

4. Bizhanova, A., and Kaufman, P.D. (2021). Close to the edge: Heterochromatin at the nucleolar and nuclear peripheries. Biochim Biophys Acta Gene Regul Mech 1864, 194666. 10.1016/j.bbagrm.2020.194666.

5. Briand, N., and Collas, P. (2018). Laminopathy-causing lamin A mutations reconfigure lamina-associated domains and local spatial chromatin conformation. Nucleus 9, 216–226. 10.1080/19491034.2018.1449498.

6. Guelen, L., Pagie, L., Brasset, E., Meuleman, W., Faza, M.B., Talhout, W., Eussen, B.H., de Klein, A., Wessels, L., de Laat, W., and van Steensel, B. (2008). Domain organization of human chromosomes revealed by mapping of nuclear lamina interactions. Nature 453, 948–951. 10.1038/nature06947.

7. Peric-Hupkes, D., Meuleman, W., Pagie, L., Bruggeman, S.W., Solovei, I., Brugman, W., Graf, S., Flicek, P., Kerkhoven, R.M., van Lohuizen, M., et al. (2010). Molecular maps of the reorganization of genome-nuclear lamina interactions during differentiation. Mol Cell 38, 603–613. 10.1016/j.molcel.2010.03.016.

8. Ahanger, S.H., Delgado, R.N., Gil, E., Cole, M.A., Zhao, J., Hong, S.J., Kriegstein, A.R., Nowakowski, T.J., Pollen, A.A., and Lim, D.A. (2021). Distinct nuclear compartment-associated genome architecture in the developing mammalian brain. Nat Neurosci 24, 1235–1242. 10.1038/s41593-021-00879-5.

9. Meuleman, W., Peric-Hupkes, D., Kind, J., Beaudry, J.B., Pagie, L., Kellis, M., Reinders, M., Wessels, L., and van Steensel, B. (2013). Constitutive nuclear lamina-genome interactions are highly conserved and associated with A/T-rich sequence. Genome Res 23, 270–280. 10.1101/gr.141028.112.

10. Falk, M., Feodorova, Y., Naumova, N., Imakaev, M., Lajoie, B.R., Leonhardt, H., Joffe, B., Dekker, J., Fudenberg, G., Solovei, I., and Mirny, L.A. (2019). Heterochromatin drives compartmentalization of inverted and conventional nuclei. Nature 570, 395–399. 10.1038/s41586-019-1275-3.

11. Bajpai, G., Amiad Pavlov, D., Lorber, D., Volk, T., and Safran, S. (2021). Mesoscale phase separation of chromatin in the nucleus. Elife 10. 10.7554/eLife.63976.

12. Maji, A., Ahmed, J.A., Roy, S., Chakrabarti, B., and Mitra, M.K. (2020). A Lamin-Associated Chromatin Model for Chromosome Organization. Biophys J 118, 3041–3050. 10.1016/j.bpj.2020.05.014.

13. Chiang, M., Michieletto, D., Brackley, C.A., Rattanavirotkul, N., Mohammed, H., Marenduzzo, D., and Chandra, T. (2019). Polymer Modeling Predicts Chromosome Reorganization in Senescence. Cell Rep 28, 3212–3223 e3216. 10.1016/j.celrep.2019.08.045.

14. Amiad-Pavlov, D., Lorber, D., Bajpai, G., Reuveny, A., Roncato, F., Alon, R., Safran, S., and Volk, T. (2021). Live imaging of chromatin distribution reveals novel principles of nuclear architecture and chromatin compartmentalization. Sci Adv 7. 10.1126/sciadv.abf6251.

15. Rooijers, K., Markodimitraki, C.M., Rang, F.J., de Vries, S.S., Chialastri, A., de Luca, K.L., Mooijman, D., Dey, S.S., and Kind, J. (2019). Simultaneous quantification of protein- DNA contacts and transcriptomes in single cells. Nat Biotechnol 37, 766–772. 10.1038/s41587-019-0150-y.

16. Poleshko, A., Shah, P.P., Gupta, M., Babu, A., Morley, M.P., Manderfield, L.J., Iãovits, J.L., Calderon, D., Aghajanian, H., Sierra-Pagan, J.E., et al. (2017). Genome-Nuclear Lamina Interactions Regulate Cardiac Stem Cell Lineage Restriction. Cell 171, 573–587 e514. 10.1016/j.cell.2017.09.018.

17. Borsos, M., Perricone, S.M., Schauer, T., Pontabry, J., de Luca, K.L., de Vries, S.S., Ruiz-Morales, E.R., Torres-Padilla, M.E., and Kind, J. (2019). Genome-lamina interactions are established de novo in the early mouse embryo. Nature 569, 729–733. 10.1038/s41586-019-1233-0.

18. Finlan, L.E., Sproul, D., Thomson, I., Boyle, S., Kerr, E., Perry, P., Ylstra, B., Chubb, J.R., and Bickmore, W.A. (2008). Recruitment to the nuclear periphery can alter expression of genes in human cells. PLoS Genet 4, e1000039. 10.1371/journal.pgen.1000039.

19. Reddy, K.L., Zullo, J.M., Bertolino, E., and Singh, H. (2008). Transcriptional repression mediated by repositioning of genes to the nuclear lamina. Nature 452, 243–247. 10.1038/nature06727.

20. Kumaran, R.I., and Spector, D.L. (2008). A genetic locus targeted to the nuclear periphery in living cells maintains its transcriptional competence. J Cell Biol 180, 51–65. 10.1083/jcb.200706060.

21. Manzo, S.G., Dauban, L., and van Steensel, B. (2022). Lamina-associated domains: Tethers and looseners. Curr Opin Cell Biol 74, 80–87. 10.1016/j.ceb.2022.01.004.

22. Kiseleva, A.A., and Poleshko, A. (2023). The secret life of chromatin tethers. FEBS Leh 597, 2782–2790. 10.1002/1873-3468.14685.

23. Kind, J., Pagie, L., de Vries, S.S., Nahidiazar, L., Dey, S.S., Bienko, M., Zhan, Y., Lajoie, B., de Graaf, C.A., Amendola, M., et al. (2015). Genome-wide maps of nuclear lamina interactions in single human cells. Cell 163, 134–147. 10.1016/j.cell.2015.08.040.

24. Chubb, J.R., Boyle, S., Perry, P., and Bickmore, W.A. (2002). Chromatin motion is constrained by association with nuclear compartments in human cells. Curr Biol 12, 439–445. 10.1016/s0960-9822(02)00695-4.

25. Kind, J., Pagie, L., Ortabozkoyun, H., Boyle, S., de Vries, S.S., Janssen, H., Amendola, M., Nolen, L.D., Bickmore, W.A., and van Steensel, B. (2013). Single-cell dynamics of genome-nuclear lamina interactions. Cell 153, 178–192. 10.1016/j.cell.2013.02.028.

26. Zullo, J.M., Demarco, I.A., Pique-Regi, R., Gaffney, D.J., Epstein, C.B., Spooner, C.J., Luperchio, T.R., Bernstein, B.E., Pritchard, J.K., Reddy, K.L., and Singh, H. (2012). DNA sequence-dependent compartmentalization and silencing of chromatin at the nuclear lamina. Cell 149, 1474–1487. 10.1016/j.cell.2012.04.035.

27. Bian, Q., Khanna, N., Alvikas, J., and Belmont, A.S. (2013). beta-Globin cis-elements determine differential nuclear targeting through epigenetic modifications. J Cell Biol 203, 767–783. 10.1083/jcb.201305027.

28. Harr, J.C., Luperchio, T.R., Wong, X., Cohen, E., Wheelan, S.J., and Reddy, K.L. (2015). Directed targeting of chromatin to the nuclear lamina is mediated by chromatin state and A-type lamins. J Cell Biol 208, 33–52. 10.1083/jcb.201405110.

29. Hama, C., Ali, Z., and Kornberg, T.B. (1990). Region-specific recombination and expression are directed by portions of the Drosophila engrailed promoter. Genes Dev 4, 1079–1093. 10.1101/gad.4.7.1079.

30. Cheng, Y., Kwon, D.Y., Arai, A.L., Mucci, D., and Kassis, J.A. (2012). P-element homing is facilitated by engrailed polycomb-group response elements in Drosophila melanogaster. PLoS One 7, e30437. 10.1371/journal.pone.0030437.

31. Lieberman-Aiden, E., van Berkum, N.L., Williams, L., Imakaev, M., Ragoczy, T., Telling, A., Amit, I., Lajoie, B.R., Sabo, P.J., Dorschner, M.O., et al. (2009). Comprehensive mapping of long-range interactions reveals folding principles of the human genome. Science 326, 289–293. 10.1126/science.1181369.

32. Paulsen, J., Liyakat Ali, T.M., Nekrasov, M., Delbarre, E., Baudement, M.O., Kurscheid, S., Tremethick, D., and Collas, P. (2019). Long-range interactions between topologically associating domains shape the four-dimensional genome during differentiation. Nat Genet 51, 835–843. 10.1038/s41588-019-0392-0.

33. Kokubu, C., Horie, K., Abe, K., Ikeda, R., Mizuno, S., Uno, Y., Ogiwara, S., Ohtsuka, M., Isotani, A., Okabe, M., et al. (2009). A transposon-based chromosomal engineering method to survey a large cis-regulatory landscape in mice. Nat Genet 41, 946–952. 10.1038/ng.397.

34. Ruf, S., Symmons, O., Uslu, V.V., Dolle, D., Hot, C., Ettwiller, L., and Spitz, F. (2011). Largescale analysis of the regulatory architecture of the mouse genome with a transposon-associated sensor. Nat Genet 43, 379–386. 10.1038/ng.790.

35. Shaw, D., Miravet-Verde, S., Pinero-Lambea, C., Serrano, L., and Lluch-Senar, M. (2021). LoxTnSeq: random transposon insertions combined with cre/lox recombination and counterselection to generate large random genome reductions. Microb Biotechnol 14, 2403–2419. 10.1111/1751-7915.13714.

36. Liang, Q., Kong, J., Stalker, J., and Bradley, A. (2009). Chromosomal mobilization and reintegration of Sleeping Beauty and PiggyBac transposons. Genesis 47, 404–408. 10.1002/dvg.20508.

37. Yoshida, J., Akagi, K., Misawa, R., Kokubu, C., Takeda, J., and Horie, K. (2017). Chromatin states shape insertion profiles of the piggyBac, Tol2 and Sleeping Beauty transposons and murine leukemia virus. Sci Rep 7, 43613. 10.1038/srep43613.

38. Yant, S.R., Wu, X., Huang, Y., Garrison, B., Burgess, S.M., and Kay, M.A. (2005). High-resolution genome-wide mapping of transposon integration in mammals. Mol Cell Biol 25, 2085–2094. 10.1128/MCB.25.6.2085-2094.2005.

39. de Jong, J., Akhtar, W., Badhai, J., Rust, A.G., Rad, R., Hilkens, J., Berns, A., van Lohuizen, M., Wessels, L.F., and de Ridder, J. (2014). Chromatin landscapes of retroviral and transposon integration profiles. PLoS Genet 10, e1004250. 10.1371/journal.pgen.1004250.

40. Luo, G., Ivics, Z., Izsvak, Z., and Bradley, A. (1998). Chromosomal transposition of a Tc1/mariner-like element in mouse embryonic stem cells. Proc Natl Acad Sci U S A 95, 10769–10773. 10.1073/pnas.95.18.10769.

41. Brueckner, L., Zhao, P.A., van Schaik, T., Leemans, C., Sima, J., Peric-Hupkes, D., Gilbert, D.M., and van Steensel, B. (2020). Local rewiring of genome-nuclear lamina interactions by transcription. EMBO J 39, e103159. 10.15252/embj.2019103159.

42. Jin, Z., Maiti, S., Huls, H., Singh, H., Olivares, S., Mates, L., Izsvak, Z., Ivics, Z., Lee, D.A., Champlin, R.E., and Cooper, L.J. (2011). The hyperactive Sleeping Beauty transposase SB100X improves the genetic modification of T cells to express a chimeric antigen receptor. Gene Ther 18, 849–856. 10.1038/gt.2011.40.

43. Stern, D.L. (2017). Tagmentation-Based Mapping (TagMap) of Mobile DNA Genomic Insertion Sites. bioRxiv, 037762. 10.1101/037762.

44. Zheng, X., Hu, J., Yue, S., Kristiani, L., Kim, M., Sauria, M., Taylor, J., Kim, Y., and Zheng, Y. (2018). Lamins Organize the Global Three-Dimensional Genome from the Nuclear Periphery. Mol Cell 71, 802–815 e807. 10.1016/j.molcel.2018.05.017.

45. van Schaik, T., Vos, M., Peric-Hupkes, D., Hn Celie, P., and van Steensel, B. (2020). Cell cycle dynamics of lamina-associated DNA. EMBO Rep 21, e50636. 10.15252/embr.202050636.

46. Jinek, M., East, A., Cheng, A., Lin, S., Ma, E., and Doudna, J. (2013). RNA-programmed genome editing in human cells. Elife 2, e00471. 10.7554/eLife.00471.

47. Cong, L., Ran, F.A., Cox, D., Lin, S., Barretto, R., Habib, N., Hsu, P.D., Wu, X., Jiang, W., Marraffini, L.A., and Zhang, F. (2013). Multiplex genome engineering using CRISPR/Cas systems. Science 339, 819–823. 10.1126/science.1231143.

48. Mali, P., Yang, L., Esvelt, K.M., Aach, J., Guell, M., DiCarlo, J.E., Norville, J.E., and Church, G.M. (2013). RNA-guided human genome engineering via Cas9. Science 339, 823–826. 10.1126/science.1232033.

49. Vorontsov, I.E., Eliseeva, I.A., Zinkevich, A., Nikonov, M., Abramov, S., Boytsov, A., Kamenets, V., Kasianova, A., Kolmykov, S., Yevshin, I.S., et al. (2023). HOCOMOCO in 2024: a rebuild of the curated collection of binding models for human and mouse transcription factors. Nucleic Acids Res. 10.1093/nar/gkad1077.

50. Lu, J.Y., Shao, W., Chang, L., Yin, Y., Li, T., Zhang, H., Hong, Y., Percharde, M., Guo, L., Wu, Z., et al. (2020). Genomic Repeats Categorize Genes with Distinct Functions for Orchestrated Regulation. Cell Rep 30, 3296–3311 e3295. 10.1016/j.celrep.2020.02.048.

51. Vazquez, B.N., Thackray, J.K., Simonet, N.G., Chahar, S., Kane-Goldsmith, N., Newkirk, S.J., Lee, S., Xing, J., Verzi, M.P., An, W., et al. (2019). SIRT7 mediates L1 elements transcriptional repression and their association with the nuclear lamina. Nucleic Acids Res 47, 7870–7885. 10.1093/nar/gkz519.

52. Dixon, J.R., Selvaraj, S., Yue, F., Kim, A., Li, Y., Shen, Y., Hu, M., Liu, J.S., and Ren, B. (2012). Topological domains in mammalian genomes identified by analysis of chromatin interactions. Nature 485, 376–380. 10.1038/nature11082.

53. Rao, S.S., Huntley, M.H., Durand, N.C., Stamenova, E.K., Bochkov, I.D., Robinson, J.T., Sanborn, A.L., Machol, I., Omer, A.D., Lander, E.S., and Aiden, E.L. (2014). A 3D map of the human genome at kilobase resolution reveals principles of chromatin looping. Cell 159, 1665–1680. 10.1016/j.cell.2014.11.021.

54. Spracklin, G., Abdennur, N., Imakaev, M., Chowdhury, N., Pradhan, S., Mirny, L.A., and Dekker, J. (2023). Diverse silent chromatin states modulate genome compartmentalization and loop extrusion barriers. Nat Struct Mol Biol 30, 38–51. 10.1038/s41594-022-00892-7.

55. Towbin, B.D., Gonzalez-Aguilera, C., Sack, R., Gaidatzis, D., Kalck, V., Meister, P., Askjaer, P., and Gasser, S.M. (2012). Step-wise methylation of histone H3K9 positions heterochromatin at the nuclear periphery. Cell 150, 934–947. 10.1016/j.cell.2012.06.051.

56. Wu, F., and Yao, J. (2017). Identifying Novel Transcriptional and Epigenetic Features of Nuclear Lamina-associated Genes. Sci Rep 7, 100. 10.1038/s41598-017-00176-x.

57. Leemans, C., van der Zwalm, M.C.H., Brueckner, L., Comoglio, F., van Schaik, T., Pagie, L., van Arensbergen, J., and van Steensel, B. (2019). Promoter-Intrinsic and Local Chromatin Features Determine Gene Repression in LADs. Cell 177, 852–864 e814. 10.1016/j.cell.2019.03.009.

58. Brunet, A., Destainville, N., and Collas, P. (2021). Physical constraints in polymer modeling of chromatin associations with the nuclear periphery at kilobase scale. Nucleus 12, 6–20. 10.1080/19491034.2020.1868105.

59. Tortora, M.M.C., Brennan, L.D., Karpen, G., and Jost, D. (2023). HP1-driven phase separation recapitulates the thermodynamics and kinetics of heterochromatin condensate formation. Proc Natl Acad Sci U S A 120, e2211855120. 10.1073/pnas.2211855120.

60. Therizols, P., Illingworth, R.S., Courilleau, C., Boyle, S., Wood, A.J., and Bickmore, W.A. (2014). Chromatin decondensation is sufficient to alter nuclear organization in embryonic stem cells. Science 346, 1238–1242. 10.1126/science.1259587.

61. Chen, S., Luperchio, T.R., Wong, X., Doan, E.B., Byrd, A.T., Roy Choudhury, K., Reddy, K.L., and Krangel, M.S. (2018). A Lamina-Associated Domain Border Governs Nuclear Lamina Interactions, Transcription, and Recombination of the Tcrb Locus. Cell Rep 25, 1729–1740 e1726. 10.1016/j.celrep.2018.10.052.

62. Isoda, T., Moore, A.J., He, Z., Chandra, V., Aida, M., Denholtz, M., Piet van Hamburg, J., Fisch, K.M., Chang, A.N., Fahl, S.P., et al. (2017). Non-coding Transcription Instructs Chromatin Folding and Compartmentalization to Dictate Enhancer-Promoter Communication and T Cell Fate. Cell 171, 103–119 e118. 10.1016/j.cell.2017.09.001.

63. Clowney, E.J., LeGros, M.A., Mosley, C.P., Clowney, F.G., Markenskoff-Papadimitriou, E.C., Myllys, M., Barnea, G., Larabell, C.A., and Lomvardas, S. (2012). Nuclear aggregation of olfactory receptor genes governs their monogenic expression. Cell 151, 724–737. 10.1016/j.cell.2012.09.043.

64. Solovei, I., Wang, A.S., Thanisch, K., Schmidt, C.S., Krebs, S., Zwerger, M., Cohen, T.V., Devys, D., Foisner, R., Peichl, L., et al. (2013). LBR and lamin A/C sequentially tether peripheral heterochromatin and inversely regulate differentiation. Cell 152, 584–598. 10.1016/j.cell.2013.01.009.

65. Duband-Goulet, I., and Courvalin, J.C. (2000). Inner nuclear membrane protein LBR preferentially interacts with DNA secondary structures and nucleosomal linker. Biochemistry 39, 6483–6488. 10.1021/bi992908b.

66. Hoskins, V.E., Smith, K., and Reddy, K.L. (2021). The shifting shape of genomes: dynamics of heterochromatin interactions at the nuclear lamina. Curr Opin Genet Dev 67, 163–173. 10.1016/j.gde.2021.02.003.

67. Robson, M.I., de Las Heras, J.I., Czapiewski, R., Le Thanh, P., Booth, D.G., Kelly, D.A., Webb, S., Kerr, A.R.W., and Schirmer, E.C. (2016). Tissue-Specific Gene Repositioning by Muscle Nuclear Membrane Proteins Enhances Repression of Critical Developmental Genes during Myogenesis. Mol Cell 62, 834–847. 10.1016/j.molcel.2016.04.035.

68. Rivera-Mulia, J.C., Dimond, A., Vera, D., Trevilla-Garcia, C., Sasaki, T., Zimmerman, J., Dupont, C., Gribnau, J., Fraser, P., and Gilbert, D.M. (2018). Allele-specific control of replication timing and genome organization during development. Genome Res 28, 800–811. 10.1101/gr.232561.117.

69. Labun, K., Montague, T.G., Krause, M., Torres Cleuren, Y.N., Tjeldnes, H., and Valen, E. (2019). CHOPCHOP v3: expanding the CRISPR web toolbox beyond genome editing. Nucleic Acids Res 47, W171–W174. 10.1093/nar/gkz365.

70. Montague, T.G., Cruz, J.M., Gagnon, J.A., Church, G.M., and Valen, E. (2014). CHOPCHOP: a CRISPR/Cas9 and TALEN web tool for genome editing. Nucleic Acids Res 42, W401–407. 10.1093/nar/gku410.

71. Brinkman, E.K., Chen, T., Amendola, M., and van Steensel, B. (2014). Easy quantitative assessment of genome editing by sequence trace decomposition. Nucleic Acids Res 42, e168. 10.1093/nar/gku936.

72. Schep, R., Brinkman, E.K., Leemans, C., Vergara, X., van der Weide, R.H., Morris, B., van Schaik, T., Manzo, S.G., Peric-Hupkes, D., van den Berg, J., et al. (2021). Impact of chromatin context on Cas9-induced DNA double-strand break repair pathway balance. Mol Cell 81, 2216–2230 e2210. 10.1016/j.molcel.2021.03.032.

73. Manjon, A.G., Manzo, S.G., Prekovic, S., Potgeter, L., van Schaik, T., Liu, N.Q., Flach, K., Peric-Hupkes, D., Joosten, S., Teunissen, H., et al. (2023). Perturbations in 3D genome organization can promote acquired drug resistance. Cell Rep 42, 113124. 10.1016/j.celrep.2023.113124.

74. Zuin, J., Roth, G., Zhan, Y., Cramard, J., Redolfi, J., Piskadlo, E., Mach, P., Kryzhanovska, M., Tihanyi, G., Kohler, H., et al. (2022). Nonlinear control of transcription through enhancer-promoter interactions. Nature 604, 571–577. 10.1038/s41586-022-04570-y.

75. Frankish, A., Diekhans, M., Ferreira, A.M., Johnson, R., Jungreis, I., Loveland, J., Mudge, J.M., Sisu, C., Wright, J., Armstrong, J., et al. (2019). GENCODE reference annotation for the human and mouse genomes. Nucleic Acids Res 47, D766–D773. 10.1093/nar/gky955.

76. van de Geijn, B., McVicker, G., Gilad, Y., and Pritchard, J.K. (2015). WASP: allele-specific software for robust molecular quantitative trait locus discovery. Nat Methods 12, 1061–1063. 10.1038/nmeth.3582.

77. Keane, T.M., Goodstadt, L., Danecek, P., White, M.A., Wong, K., Yalcin, B., Heger, A., Agam, A., Slater, G., Goodson, M., et al. (2011). Mouse genomic variation and its effect on phenotypes and gene regulation. Nature 477, 289–294. 10.1038/nature10413.

78. Doran, A.G., Wong, K., Flint, J., Adams, D.J., Hunter, K.W., and Keane, T.M. (2016). Deep genome sequencing and variation analysis of 13 inbred mouse strains defines candidate phenotypic alleles, private variation and homozygous truncating mutations. Genome Biol 17, 167. 10.1186/s13059-016-1024-y.

79. Fornes, O., Castro-Mondragon, J.A., Khan, A., van der Lee, R., Zhang, X., Richmond, P.A., Modi, B.P., Correard, S., Gheorghe, M., Baranasic, D., et al. (2020). JASPAR 2020: update of the open-access database of transcription factor binding profiles. Nucleic Acids Res 48, D87–D92. 10.1093/nar/gkz1001.

80. Dobin, A., Davis, C.A., Schlesinger, F., Drenkow, J., Zaleski, C., Jha, S., Batut, P., Chaisson, M., and Gingeras, T.R. (2013). STAR: ultrafast universal RNA-seq aligner. Bioinformatics 29, 15–21. 10.1093/bioinformatics/bts635.

81. Li, H., Handsaker, B., Wysoker, A., Fennell, T., Ruan, J., Homer, N., Marth, G., Abecasis, G., Durbin, R., and Genome Project Data Processing, S. (2009). The Sequence Alignment/Map format and SAMtools. Bioinformatics 25, 2078-2079. 10.1093/bioinformatics/btp352.

82. Shumate, A., and Salzberg, S.L. (2021). Liftoff: accurate mapping of gene annotations. Bioinformatics 37, 1639–1643. 10.1093/bioinformatics/btaa1016.

83. Lawrence, M., Huber, W., Pages, H., Aboyoun, P., Carlson, M., Gentleman, R., Morgan, M.T., and Carey, V.J. (2013). Software for computing and annotating genomic ranges. PLoS Comput Biol 9, e1003118. 10.1371/journal.pcbi.1003118.

